# The nucleocytosolic *O*-fucosyltransferase *Spindly* affects protein expression and virulence in *Toxoplasma gondii*

**DOI:** 10.1101/2020.08.30.274134

**Authors:** Giulia Bandini, Carolina Agop-Nersesian, Hanke van der Wel, Msano Mandalasi, Hyunwoo Kim, Christopher M. West, John Samuelson

## Abstract

Once considered unusual, nucleocytoplasmic glycosylation is now recognized as a conserved feature of eukaryotes. While in animals *O*-GlcNAc transferase (OGT) modifies thousands of intracellular proteins, the human pathogen *Toxoplasma gondii* transfers a different sugar, fucose, to proteins involved in transcription, mRNA processing and signaling. Knockout experiments showed that *Tg*SPY, an ortholog of plant SPINDLY and paralog of host OGT, is required for nuclear *O*-fucosylation. Here we verify that *Tg*SPY is the nucleocytoplasmic *O*-fucosyltransferase (OFT) by 1) complementation with *Tg*SPY-MYC_3_, 2) its functional dependence on amino acids critical for OGT activity, and 3) its ability to *O*-fucosylate itself and a model substrate and to specifically hydrolyze GDP-Fuc. While many of the endogenous proteins modified by *O*-Fuc are important for tachyzoite fitness, *O*-fucosylation by *Tg*SPY is not essential. Growth of Δ*spy* tachyzoites in fibroblasts is modestly affected, despite marked reductions in the levels of ectopically-expressed proteins normally modified with *O*-fucose. Intact *Tg*SPY-MYC_3_ localizes to the nucleus and cytoplasm, whereas catalytic mutants often displayed reduced abundance. Δ*spy* tachyzoites of a luciferase-expressing type II strain exhibited infection kinetics in mice similar to wild type but increased persistence in the chronic brain phase, potentially due to an imbalance of regulatory protein levels. The modest changes in parasite fitness *in vitro* and in mice, despite profound effects on reporter protein accumulation, and the characteristic punctate localization of *O*-fucosylated proteins, suggest that *Tg*SPY controls the levels of proteins to be held in reserve for response to novel stresses.

## Introduction

*Toxoplasma gondii* is an obligate intracellular protist classified as a Category B pathogen on NIAID’s list of emerging infectious diseases (1). Based on serological evidence, the CDC estimates that *T. gondii* has infected ∼11% of the United States population, while infection rates in other countries may exceed 50% (2, 3). Humans can become infected with *T. gondii* when they ingest food contaminated with oocysts shed in the feces of cats (where the sexual cycle occurs) or tissue cysts present in undercooked meat (4). *T. gondii* infection is generally transient and presents with no or mild symptoms, except in immunocompromised individuals and in fetuses; in these cases severe ocular and neurologic disease may occur (5, 6). Because it is difficult to prevent *T. gondii* infection, congenital transmission remains a significant problem in the US and around the world, together with encephalitis caused by reactivation of the parasite in immunosuppressed individuals (7, 8). Even though prior exposure is believed to lead to immunity against reinfection, significant challenges persist: there is no vaccine against *T. gondii* (9), current drug treatments are effective only in acute infections, and development of drug resistance remains a concern (10).

*T. gondii* goes through a series of differentiation events in both its asexual and sexual cycles (1). During asexual development in humans and other intermediate hosts, tachyzoites (the fast replicative form of the parasite) differentiate to bradyzoites (the slow replicative form) when exposed to unfavorable conditions, *e*.*g*. immune system response or reduced availability of nutrients (11). *T. gondii* bradyzoites have a slower metabolic rate and organize in cysts, which are characterized by a wall that is rich in glycoconjugates and are found predominantly in muscles and in the brain (12, 13). Importantly, both drug treatment failure and latency are linked to parasite persistence in tissue cysts (11).

Transcriptomics and proteomic studies of developmental stages of *T. gondii* have identified many stage-specific genes and proteins, indicating the importance of tightly regulated expression patterns (14, 15). In support of this idea, master regulators of bradyzoites differentiation (BDF1) and sexual commitment (MORC) have been recently identified (13, 16). Many of the downstream transcriptional and a few translational factors in the parasite have been shown to be regulated by post-translational modifications (PTMs), including phosphorylation, acetylation and ubiquitination (17–20). While transcriptional regulation of protein expression has been the subject of many studies in *T. gondii*, the role that PTMs might play in regulating protein stability, concentration, and/or localization (protein homeostasis) are less understood.

Our recent studies identified *O*-linked fucose (*O*-Fuc) as an additional PTM that modifies *T. gondii* proteins that are involved in regulation of protein expression and cell signaling. We showed that the L-fucose-specific lectin from *Aleuria aurantia* (AAL) highlights the nucleoplasm and in particular the nuclear periphery of RH, a *T. gondii* type I strain, but not the secretory system, in contrast to what is observed in most eukaryotic cells (21, 22). Super-resolution microscopy showed that *O*-fucosylated proteins localize in a punctate pattern in the vicinity of nuclear pore complexes (NPCs), suggesting they might be forming assemblies. Mass spectrometry of parasite AAL-enriched proteins revealed a high incidence of one or more fucose residues *O*-linked to Ser/Thr on peptides from intrinsically disordered regions (IDRs) on 69 proteins, including Phe/Gly-repeats nucleoporins (FG-Nups), mRNA processing enzymes such as members of the CCR4-NOT complex, transcriptional regulators such as a subunit of the transcription initiation complex, and cell-signaling proteins such as kinases and ubiquitination factors (23). A yellow fluorescent protein-fusion containing a Ser-rich domain (SRD-YFP) from a *T. gondii* GPN-loop GTPase (*Tg*GPN), which is 79-amino acids long and contains 37 consecutive Ser residues, was found to be modified with *O*-Fuc and partially associated with AAL-labeled assemblies, while fucosylation was not detected on YFP with a nuclear localization signal (NLS) (22). The proteins and sequences modified with *O*-Fuc in *T. gondii* closely resemble the substrates of the human OGT (*Hs*OGT). *O*-GlcNAcylation is important in disease-relevant signaling and enzyme regulation and is the best-studied example of intracellular glycosylation (24, 25).

Subsequent to the discovery of *O*-fucosylated proteins in the nucleus of *T. gondii*, the *Arabidopsis thaliana* SPINDLY (*At*SPY), a paralog of animal nucleocytoplasmic *O*-GlcNAc transferases (OGTs), was shown to be an *O*-fucosyltransferase (OFT) that activates the nuclear growth repressor DELLA (26–28). OGTs, *At*SPY, and its *T. gondii* orthologue (*Tg*SPY) each have a C-terminal Carbohydrate Active EnZyme (CAZy) glycosyltransferase 41 (GT41) catalytic domain (29) and N-terminal tetratricopeptide repeats (TPRs), 13.5 repeats in *Hs*OGT and 11 in both *At*SPY and *Tg*SPY (30). CRISPR/Cas9-mediated knockout of *Tg*SPY resulted in a modest growth phenotype, despite complete loss of AAL labeling of nuclei and parasite extracts (31). These results, pointing to OFT activity of both *Tg*SPY and *At*SPY, were quite unexpected because of their sequence homology to OGTs and because members of the same CAZy GT family usually utilize activated sugar donors with a conserved nucleotide moiety, *e*.*g*. either UDP- or GDP-sugars (29).

To better understand the cell and molecular biology of *O*-fucosylation in *T. gondii*, we tested the activity of *Tg*SPY either produced as a recombinant enzyme in bacteria or expressed in the Δ*spy* strain, with and without point mutations known to affect either host OGT or *At*SPY activities. We determined the effect of the *Tg*SPY knockout on the stability and localization of proteins bearing domains normally modified with *O*-Fuc and on the growth and differentiation of Δ*spy* strains in culture and in mice.

## Results

### The OFT activity of His-tagged *Tg*SPY (*Tg*His_6_SPY) purified from the cytosol of bacteria was shown in four ways

First, using GDP-Glo and UDP-Glo assays, in which hydrolysis of the nucleotide-sugar produces a luminescent signal, we showed that *Tg*His_6_SPY (Fig. S1), in the absence of an acceptor substrate, hydrolyzed GDP-Fuc but not UDP-GlcNAc (donor for OGT), GDP-Man, or UDP-Gal (Fig. 1A) (32). As a positive control, UDP-Gal was hydrolyzed by a *T. gondii* galactosyltransferase (*Tg*Gat1), which adds a terminal galactose to the pentasaccharide attached to the *T. gondii* Skp1 E3 ubiquitin ligase subunit (33). Second, in the presence of GDP-Fuc, *Tg*His_6_SPY fucosylated itself (OFT activity in *cis*), as shown by Western blotting with AAL (Fig. 1B). As a control, AAL binding to *Tg*His_6_SPY was inhibited by α-methyl L-fucopyranoside (Me-αFuc). To confirm that Fuc became *O*-linked to the side chain of Ser or Thr, we took advantage of knowledge that such linkages are uniquely susceptible to mild base with release of the *O*-linked Fuc and its conversion to fucitol. To detect the presence of fucitol, the product mixture from an autofucosylation reaction conducted in the presence of GDP-[^3^H]Fuc was subjected to β-elimination and the released material was chromatographically analyzed by HPAEC. As shown in Fig. 1C, all detectable radioactivity co-eluted with the fucitol standard, which was detected by pulsed amperometry. This finding confirmed that radioactivity was incorporated as Fuc and strongly supported its linkage to Ser or Thr.

**Figure 1.**
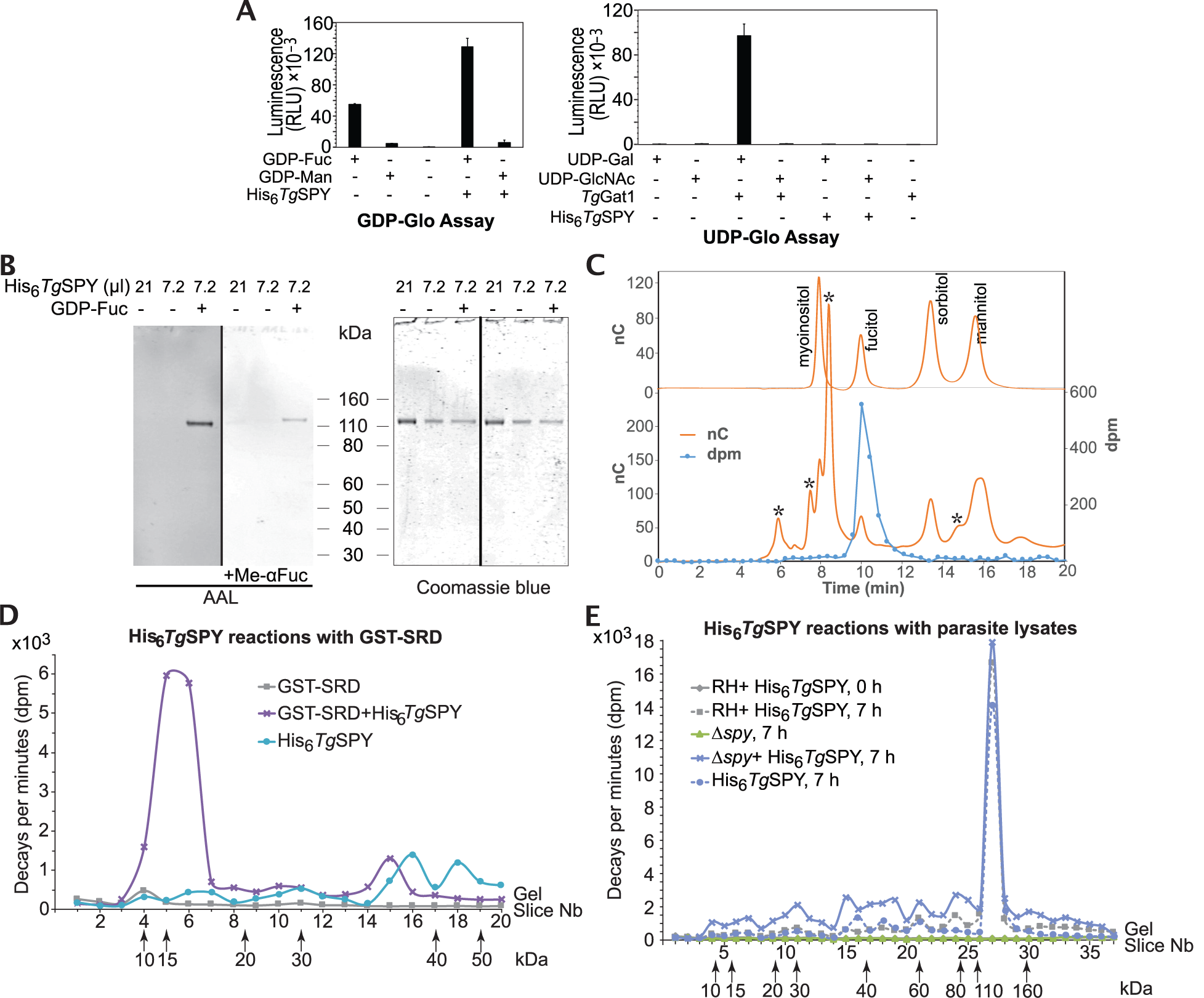
Recombinant His_6_*Tg*SPY can hydrolyze GDP-Fuc and is active against protein substrates including itself. *A*. Sugar nucleotide hydrolysis. Acceptor-independent consumption of the indicated sugar nucleotide donor substrates was assayed based on quantitation of GDP or UDP reaction products using GDP-Glo™ (upper panel) or UDP-Glo™ (lower panel) assays. Reactions were conducted in the presence or absence of either His_6_*Tg*SPY or *Tg*Gat1, an α-galactose transferase that serves as a positive control for UDP-Gal hydrolysis. The high level of GDP in the absence of enzyme is due to the intrinsic instability of GDP-Fuc. *B*. The same preparation of highly purified recombinant His_6_*Tg*SPY was incubated in the presence or absence of 2 µM GDP-Fuc. The indicated volumes were subjected to SDS-PAGE and Western blotted with biotinylated-AAL to detect fucosylation (left panel). A parallel blot was incubated with biotinylated-AAL in the presence of α-methyl fucopyranoside (Me-αFuc) as a competitive inhibitor (middle panel). The blotted gel was stained with Coomassie blue to confirm equal loading and indicated the purity of the His_6_*Tg*SPY preparation (right panel). *C*. A parallel His_6_*Tg*SPY reaction that was autofucosylated in the presence of 2 µM GDP-[^3^H]Fuc was subjected to SDS-PAGE and electroblotting onto a PVDF membrane. The SPY band was subjected to conditions of reductive β-elimination and analyzed by HPAEC on a CarboPac MA-1 column. *Top*: separation of a panel of sugar alcohols detected electrochemically. *Bottom*: separation of the reaction product spiked with the sugar alcohol sample, in a trial in which fractions were collected for analysis of radioactivity. The asterisk (*) denotes peaks of unknown origin. *D*. Purified His_6_*Tg*SPY was incubated in the presence of 2 µM GDP-[^3^H]Fuc and recombinantly prepared and purified GST-polySer as a potential acceptor substrate for 7 h. The controls omitted one or the other protein fraction. The reaction product was separated on an SDS-PAGE gel which, after Coomassie blue staining and fixation, was cut into equal sized slices and counted in a scintillation counter. *E*. Alternatively, the reaction was conducted in the presence of a desalted cytosolic extract of RH (parental) or Δ*spy* parasites for 0 or 7 h. Similar results were obtained in a complete set of independent reactions (not shown).

Third, to test for OFT activity in *trans*, we transferred the SRD of *Tg*GPN, which is *O*-fucosylated when fused to YFP in *T. gondii* tachyzoites (22), to glutathione-S-transferase (GST) to generate GST-SRD. In the presence of GDP-[^3^H]Fuc and GST-SRD, *Tg*His_6_SPY made a 15-kDa [^3^H]Fuc-labeled product, which suggested partial degradation of the substrate (Fig. 1D). Fourth, as an alternative test of OFT activity in *trans*, we incubated *Tg*His_6_SPY with desalted cytosolic extracts of Δ*spy* tachyzoites, which contain nucleocytosolic proteins that are not modified with *O*-Fuc (31). In addition to incorporating [^3^H]Fuc from GDP-[^3^H]Fuc into itself in fraction 27 (Fig. 1E), *Tg*His_6_SPY mediated substantial [^3^H]Fuc incorporation into a range of species in Δ*spy* from apparent *M*_*r*_ values of 10,000 to >200,000. Incorporation was negligible in the absence of *Tg*His_6_SPY and was substantially reduced in preparations from the wild-type RH strain, where nucleocytosolic proteins are already modified with *O*-Fuc (22).

The *in vitro* activity of *Tg*His_6_SPY strongly supports the idea that *Tg*SPY is the OFT that directly mediates assembly of the *O*-Fuc linkage in cells, as shown previously for *At*SPY (26).

### Complementation of Δ*spy* tachyzoites with myc-tagged *Tg*SPY restored *O*-Fuc modification of nucleocytosolic proteins

To further demonstrate that *Tg*SPY is the OFT, we complemented the Δ*spy* RH strain (31) with a C-terminally MYC_3_-tagged copy of *Tg*SPY cDNA, which was randomly integrated into the genome. The strategy is illustrated in Fig. 2A, and PCR validation is shown in Fig. S2A. Expression of *Tg*SPY-MYC_3_ in Δ*spy* tachyzoites cultured in human foreskin fibroblasts (HFFs) restored AAL binding to *O*-Fuc proteins of tachyzoites, as detected by structured illumination microscopy (SIM) (Fig. 2B) and lectin blotting (Fig. 2C). While there was no increase in the number of AAL-labeled assemblies in the complemented SPY knockout strain (Δ*spy*::*Tg*SPY-MYC_3_) versus wild type, there was an increase in the overall intensity of the AAL labeling (Fig. 2D). Most likely, this result can be explained by the use of the highly active tubulin promoter to express *Tg*SPY-MYC_3_ (Fig. 2A). Consistent with the unchanged number of assemblies, AAL-labeled punctae closely overlapped with YFP-tagged Nup67, which is an FG-Nup that is not modified by *O*-Fuc (Fig. 2E). Previously, we observed that AAL labeling partially overlapped with YFP-tagged Nup68, an FG-Nup that is extensively modified with *O*-Fuc (22).

**Figure 2.**
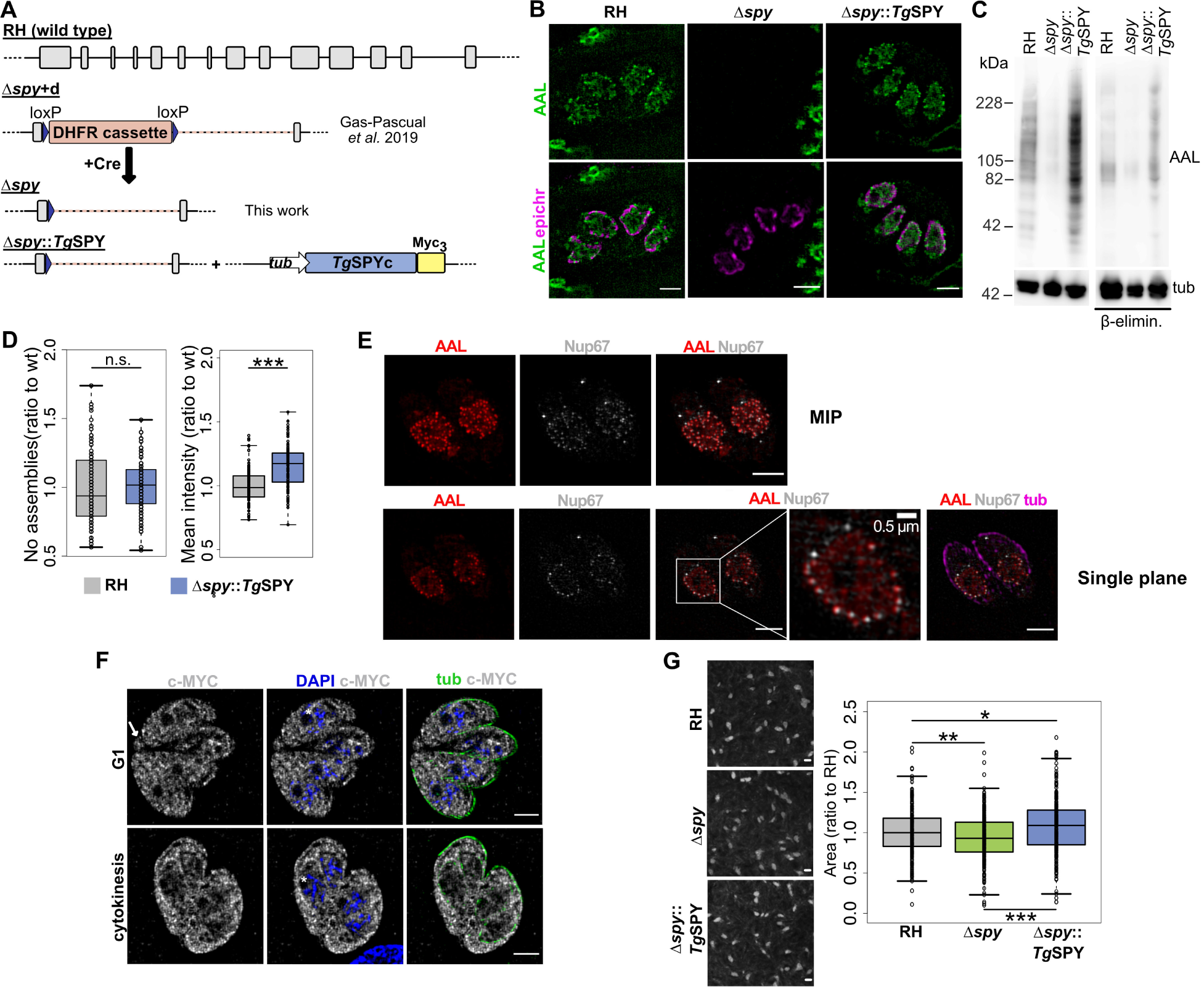
Disruption of *T. gondii spy* has a modest effect on parasite growth *in vitro* that is rescued by complementation. *A*. Schematic representation of the *spy* locus in wild type RH, the *spy* knockout, and the cell line complemented with *Tg*SPY-MYC_3_. *B*. SIM shows that nuclear staining by AAL is lost in the knockout and rescued by complementation with the endogenous enzyme. *C*. The loss and rescue of *O*-Fuc is confirmed by lectin blotting of whole cells. *D*. Quantification of the number of AAL-positive punctae (RH n=86, Δ*spy*::*Tg*SPY n=103, 3 biological repeats) and the mean intensity for the AAL signal from 3D projections (RH n=111, Δ*spy*::*Tg*SPY n=106, three biological repeats). The increase in total signal for the complemented cell line was significant (*p =* 2.6×10^-11^). *E*. SIM shows co-labeling of *O*-Fuc with AAL, tubulin with anti-tubulin, and Nup67-YFP with anti-GFP in RH parasites. The top row shows a maximum intensity projection (MIP), while the lower row shows a single plane. Nup67-YFP: an FG-Nup that has not been found to be *O*-fucosylated; epichr: epichromatin; tub: tubulin; scale bars: 2 μm, unless indicated otherwise. *F*. SIM shows that *Tg*SPY-MYC_3_ localizes to the cytosol, nucleoplasm and residual body (white arrow), but not the nucleolus (asterisks). Labeling of the parental RH strain was essentially negative at this level of exposure (data not shown). Scale bars: 0.2 cm. Box plots: the box defines the interquartile range with the black line marking the median, while the whiskers mark maximum and minimum values (excluding outliers). *G*. Plaque assays comparing wild type, knockout and complemented cell lines show that complementation with the endogenous enzyme rescues the mild growth phenotype observed in the knockout. Data are from three biological repeats. *p* values: * 0.03; ** 0.0001;*** 4×10^-11^.

*Tg*SPY-MYC_3_ was present in the cytosol, nucleoplasm, and residual body (arrow) of G1 and dividing stages of Δ*spy*::*Tg*SPY-MYC_3_ parasites (Fig. 2F). Similar to the *O*-Fuc-modified proteins visualized with AAL, *Tg*SPY-MYC_3_ was excluded from the nucleolus (asterisks). Western blot analysis of the total cell lysate from Δ*spy*::*Tg*SPY-MYC_3_ showed *Tg*SPY-MYC_3_ was full-length with the expected apparent *M*_r_ of 112,000 (Fig. 3B). Based on clonal plaque formation in fibroblast monolayers (Fig. 2G), there was a modest decrease in growth in Δ*spy* versus wild-type (RH) tachyzoites, confirming a previous report (31), and a modest increase in the Δ*spy*::*Tg*SPY-MYC_3_ strain, both of which were statistically significant.

**Figure 3.**
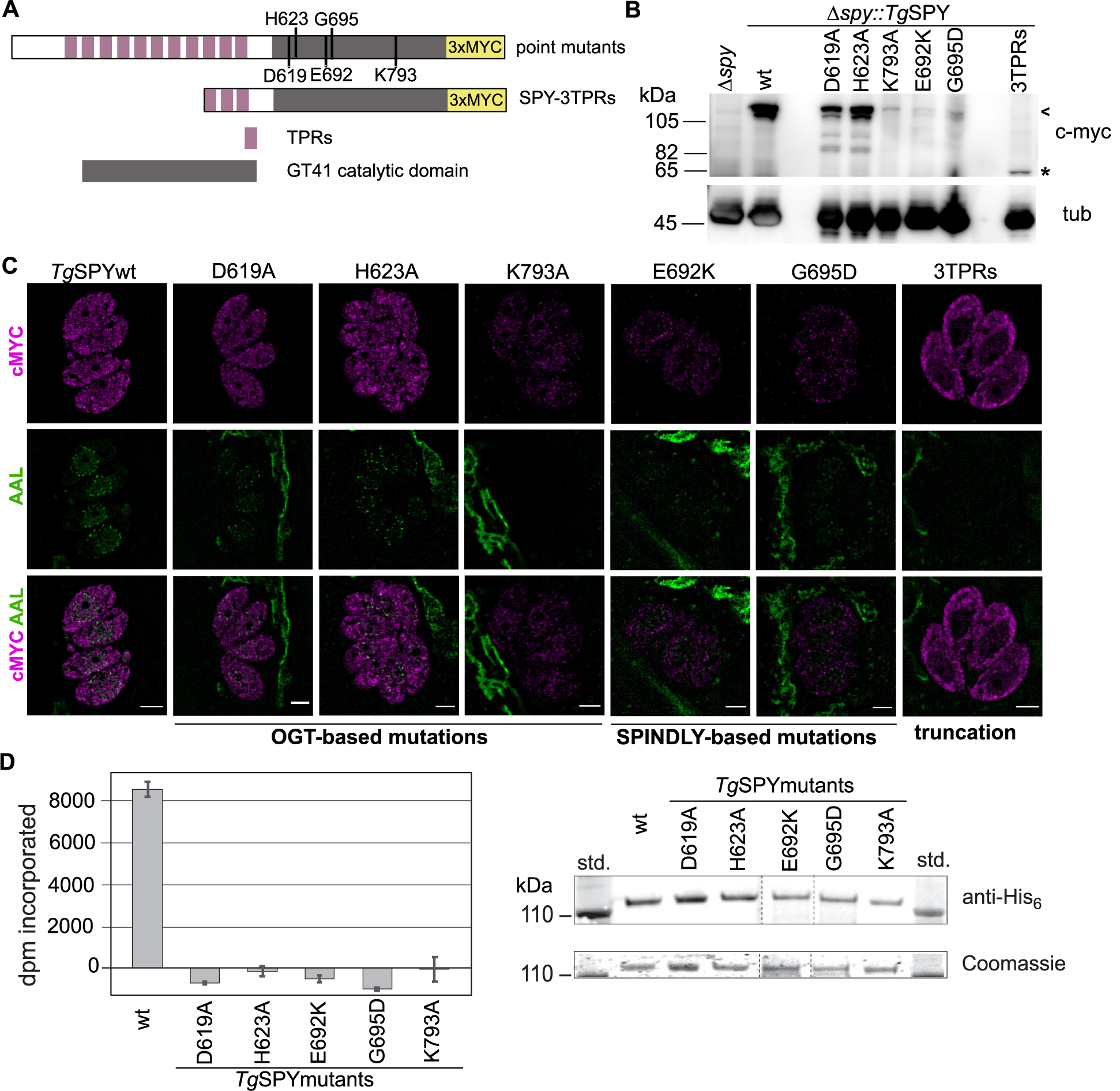
Mutagenesis analysis shows that AAL staining is dependent on the catalytic activity of *Tg*SPY and the full TPRs domain. *A*. Schematic representation of the *Tg*SPY constructs used in the mutagenesis studies. N-terminal TPRs are shown in purple and the CAZy GT41 catalytic domain in gray. The targeted residues are indicated. *B*. Western blot showing apparent *M*_*r*_ and expression levels of the mutants in the clones selected for further analysis. The position of full-length *Tg*SPY proteins is indicated by an arrowhead and that of the truncated version with an asterisk. Tubulin served as a loading control. *C*. SIM showed localization and AAL staining for all point mutants and 3TPRs truncation. Based on AAL, no activity was detected for either K793A or the truncated construct, while all other mutants showed a reduced but detectable AAL pattern. Most mutant *Tg*SPY proteins appear to be less abundant than wild type *Tg*SPY. *D*. The mutant isoforms were expressed as N-terminally His_6_-tagged proteins in *E. coli* and partially purified on Talon-Co^++^ columns. Equivalent amounts of protein were assayed for transfer of ^3^H from GDP-[^3^H]Fuc to a Ser-rich 25-mer peptide from *Tg*RINGF1. SDS-PAGE followed by Coomassie blue staining (lower right) show the calibration for the protein amounts in each reaction and was confirmed by Western blotting using anti-His_6_ (upper right). Data are from a single representative trial conducted in triplicate, and error bars represent ± standard deviation.

These complementation studies confirm that *Tg*SPY is the OFT that modifies nucleocytosolic proteins recognized by AAL and provide additional evidence that AAL-labeled assemblies co-localize with NPCs (22).

### Mutations in *Tg*SPY affected its OFT activity, its abundance, and its location

The viability of Δ*spy* parasites allowed us to investigate the biochemistry of the OFT directly in *T. gondii* tachyzoites. We generated a series of point mutations in the C-terminal GT41 catalytic domain of *Tg*SPY-MYC_3_ (D619, H623 and K793) that correspond to D554, H558 and K842 in *Hs*OGT (Fig. 3A). Mutation of either H558 or K842 to Ala abolishes glycosyltransferase activity in human and bacterial OGTs (34, 35). Structural studies show D554 is linked to its peptide substrate through a chain of water molecules, suggesting that this residue might be involved in catalysis (30). The point mutations E692K and G695D correspond to mutations in two *A. thaliana spy* hypomorphic alleles, *spy12* (G570D) and *spy15* (E567K), that have each been shown to result in complete loss or a strong reduction in OFT activity, respectively (26). Because biochemical data on *At*SPY have been obtained from experiments performed with truncated enzymes containing the C-terminal 3 TPRs (26), we also tested the behavior of a truncated *Tg*SPY-MYC_3_ containing 3 C-terminal TPRs (*Tg*SPY-3TPRs) in the Δ*spy* tachyzoites.

In Western blots of lysates of tachyzoites, all *Tg*SPY-MYC_3_ mutants were detected by anti-MYC antibodies, although the expression levels varied markedly (Fig. 3B). While D619A and H623A were present at levels comparable to the wild type, the other mutants (K793A, E692K, G695D, and the SPY-3TPRs) were present at much lower levels than wild type. By SIM, we observed that all of the point mutants of *Tg*SPY-MYC_3_, even those with very low expression levels, were located in both the cytoplasm and the nucleus, as is the case for wild-type *Tg*SPY-MYC_3_. In contrast, no truncated *Tg*SPY-3TPRs was observed in the nucleus, suggesting that the N-terminal TPRs might be involved in import of *Tg*SPY, which lacks a canonical NLS (36). AAL labeling by SIM was used to estimate *Tg*SPY activity. D619A and H623A mutants showed markedly reduced AAL labeling while only residual staining was observed in E692K and G695D. AAL labeling was absent from both the K793A mutant and the truncated SPY-3TPRs (Fig. 3C).

To confirm the occurrence of the expected negative effects of the point mutations on SPY OFT activity, N-terminally His_6_-tagged versions of the mutant isoforms were expressed in and partially purified from *E. coli*. The E692K and G695D mutants expressed with very low yield of soluble protein, and the K793A expressed at a low but better yield, which paralleled their expression levels in the parasite. A new assay that measured transfer of [^3^H]Fuc from GDP-[^3^H]Fuc to a synthetic peptide from *Tg*RINGF1 (TGGT1_285190), previously shown to be *O*-fucosylated *in vivo* (22), was developed. Relative to the wild-type protein prepared in parallel, the same concentration of each of the five point-mutants was found to exhibit no activity within the sensitivity of the assay (Fig. 3D), which is ∼10% of full activity.

We conclude that residues important for catalytic activity in *Hs*OGT and *At*SPY are also important in *Tg*SPY. Further, these point mutations appear to result in loss of protein stability in both the parasite and in *E. coli*, an effect that may be compounded by loss of potentially stabilizing auto-fucosylation. Finally, nuclear targeting of SPY appears to depend on its N-terminal TPR domains, which are also known to contribute to acceptor substrate selection and dimerization for OGT (37).

### Endogenous and exogenous proteins normally modified with *O*-Fuc were less abundant in the absence of SPY or the fucosylated domain

The goal here was to determine whether the absence of *O*-Fuc in SRDs of nucleocytosolic proteins affects their abundance and/or location. In our first experiment, we attempted to express our previously described SRD-YFP construct, which contains the 79-amino acid-long SRD from the N-terminus of *Tg*GPN that is *O*-fucosylated in wild type cells, in Δ*spy* tachyzoites. For comparison, we also expressed in Δ*spy* tachyzoites NLS-YFP, which contains a nuclear localization sequence and is not *O*-fucosylated (22). Although we were able to generate stable parasite lines expressing NLS-YFP, we could not obtain stable lines expressing SRD-YFP in Δ*spy* tachyzoites. SIM analysis of SRD-YFP was, therefore, performed by fixing fibroblast monolayers containing parental, Δ*spy* and Δ*spy*::*Tg*SPY parasites 24 h after electroporation (Fig. 4A). SRD-YFP fluorescence was strongly reduced in Δ*spy* parasites and restored in Δ*spy*::*Tg*SPY parasites. The residual labeling in Δ*spy* parasites occurred in the cytoplasm and in perinuclear puncta, in contrast to the normal enrichment in the nucleus. NLS-YFP fluorescence was comparable between parental and Δ*spy* parasites by SIM (Fig. 4B) and Western blotting (not shown). These results suggested that the SRD destabilizes YFP unless it is modified with *O*-Fuc.

**Figure 4.**
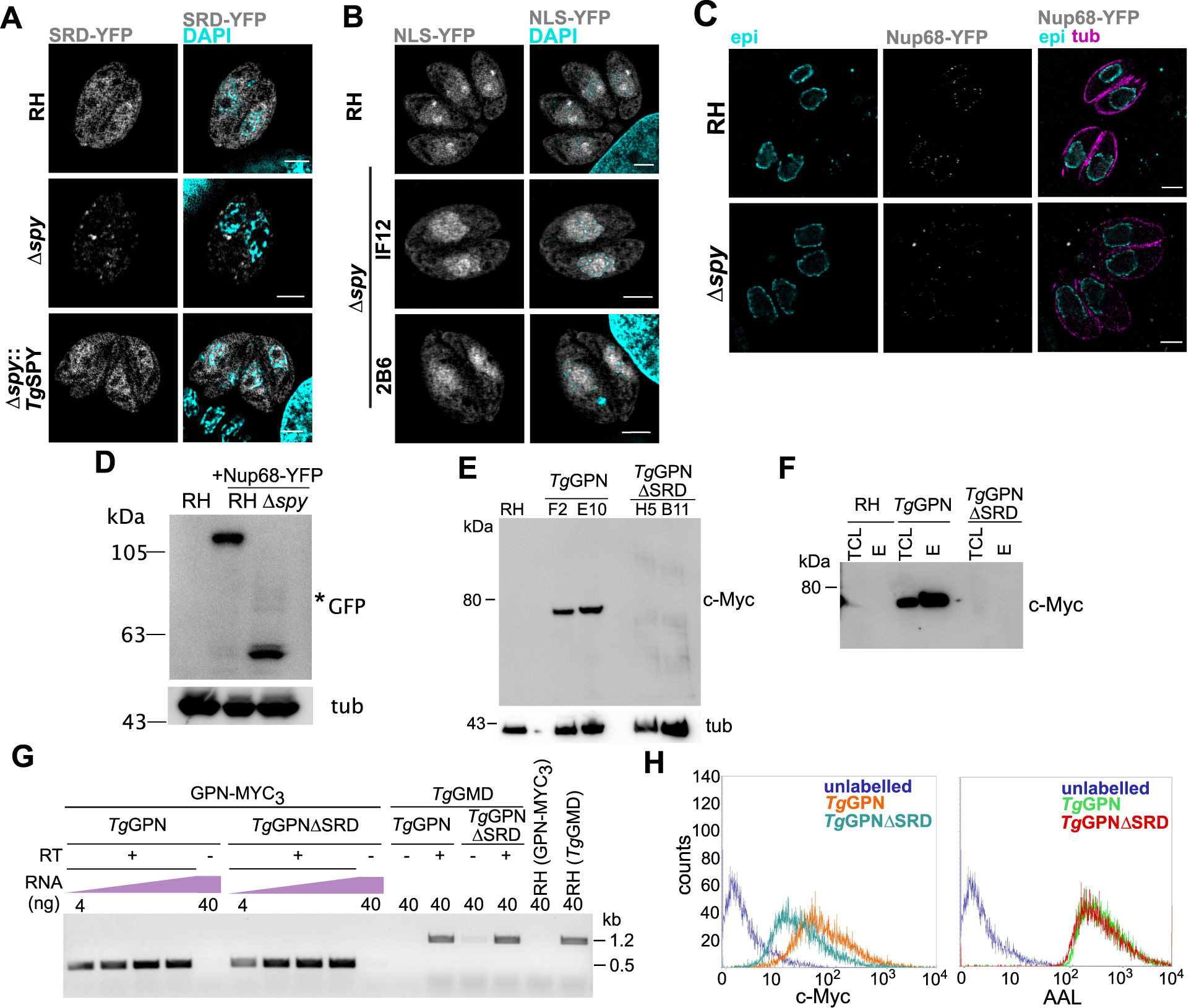
*O*-fucosylated reporter proteins stability is affected by either deletion of SRD or lack of *O*-Fuc. *A*. RH, Δ*spy* and Δ*spy*::*Tg*SPY parasites were electroporated with SRD-YFP, and expression was examined 24 h later by SIM. SRD-YFP was markedly reduced in Δ*spy* versus parental strain or complemented parasites (Δ*spy*::*Tg*SPY). *B*. A control chimera, NLS-YFP, that is not *O*-fucosylated in wild type, was stably expressed in RH, and two Δ*spy* clones showed no decrease in NLS-YFP expression versus parental strain. *C*. SIM shows Nup68-YFP, which was stably expressed under a tubulin promoter, was decreased in Δ*spy* versus parental. Colocalization was performed using antibodies against epichromatin (epi) and tubulin (tub). *D*. Nup68-YFP was detected at a lower molecular weight compared to wild type based on Western blotting with anti-GFP, suggesting it is degraded. *E*. C-terminally MYC_3_-tagged *Tg*GPN and *Tg*GPNΔSRD were ectopically expressed in RH, and two pairs of clonal populations were analyzed by Western blot with anti-cMYC. The mobility of full-length *Tg*GPN-MYC_3_ suggested an apparent *M*_r_ higher than theoretical (46,600), possibly due to the *O*-fucosylation. No clear band was observed for *Tg*GPNΔSRD. *F*. When AAL-enriched proteins were probed with anti-cMYC, full length *Tg*GPN was detected in both total cell lysate (TCL) and enriched fraction (EL), whereas but *Tg*GPNΔSRD was not detected in either fraction. *G*. Semiquantitative RT-PCR indicated comparable mRNA levels for full length and truncated isoforms. *T. gondii* GDP-mannose 4,6-dehydratase (*Tg*GMD) served as a positive control. *H*. Flow cytometry showed a decrease in mAb 9E10 binding to *Tg*GPNΔSRD-MYC_3_ compared to full length *Tg*GPN. Scale bars: 2 μm. Epi: epichromatin; tub: tubulin.

In our second experiment, we stably expressed YFP-tagged Nup68 under the tubulin promoter by random integration. In wild type RH, Nup68-YFP, which is normally heavily modified with *O*-Fuc (22), localized mainly in a punctate pattern at the nuclear periphery (Fig. 4C) and was detected by Western blot as a single band consistent with the full length fusion protein (Fig. 4D). In contrast, in Δ*spy* parasites, Nup68-YFP was barely detectable by IFA (Fig. 4C), and the protein was extensively degraded (Fig. 4D). These observations suggest that Nup68-YFP undergoes degradation in the absence of *O*-Fuc, as shown for SRD-YFP. The paucity of Nup68-YFP in Δ*spy* parasites made it impossible to judge whether the absence of *O*-Fuc affected its localization.

In a third experiment, the full-length *Tg*GPN, as well as a truncated protein missing the N-terminal SRD (*Tg*GPNΔSRD), were each tagged at the C-terminus with c-MYC_3_ and stably expressed under their own promoter after random integration into the genome (Fig. S2). In *Tg*GPNΔSRD translation was initiated at an unconserved Met residue at the beginning of the G1 motif near the start of the yeast and bacterial versions of the protein (Fig. S2D-E). Full-length *Tg*GPN accumulated at a higher *M*r than expected (49,000), based on Western blotting of two independent clones in the RH background (Fig. 4E). In contrast, *Tg*GPNΔSRD was not detected by Western blotting, even after AAL enrichment (Fig. 4F). RT-PCR demonstrated similar stable expression of the transcripts of *Tg*GPN and *Tg*GPNΔSRD (Fig. 4G), suggesting potential to express the protein. However, only a low level of expression could be detected by flow cytometry targeting the c-MYC tag (Fig. 4I). These findings are consistent with a model in which the addition of the N-terminal SRD evolved as a mechanism to allow increased translation or slower degradation kinetics of *Tg*GPN when *O*-fucosylated. It should be noted that misfolding of *Tg*GPNΔSRD due to the absence of the SRD cannot be ruled out.

### SPY modulates the growth of the type II CZ1 strain in culture and in mice

Because type I RH strain does not efficiently differentiate to bradyzoites in culture and is too virulent for mouse infection studies, we generated a *spy* knockout in a type II CZ1 Δ*ku80* strain, which was also engineered to constitutively express a firefly Re9 luciferase under control of the *gra1* promoter (see Fig. S2B and Methods for details on strain construction; Agop-Nersesian *et al*., manuscript in preparation) (38–40). Southern blot analysis was used to verify the genotype of the clonal populations (Fig. S2C). A fraction of type II strains vacuoles spontaneously differentiate to bradyzoites in normal culture conditions, as evidenced by SAG2Y expression on the parasite surface and labeling of the tissue cyst walls with *Dolichos biflorus* agglutinin (DBA), which recognizes *O*-linked *N*-acetylgalactosamine moieties on wall proteins (*e*.*g*. CST1) (12). SAG2Y-positive cells from Re9 parental strain had punctate nuclear labeling with AAL that was similar to the pattern in tachyzoites, while Δ*spy* bradyzoites had no labeling with AAL, confirming the role *Tg*SPY is required for nuclear *O*-fucosylation also in this life stage (Fig. 5A).

**Figure 5.**
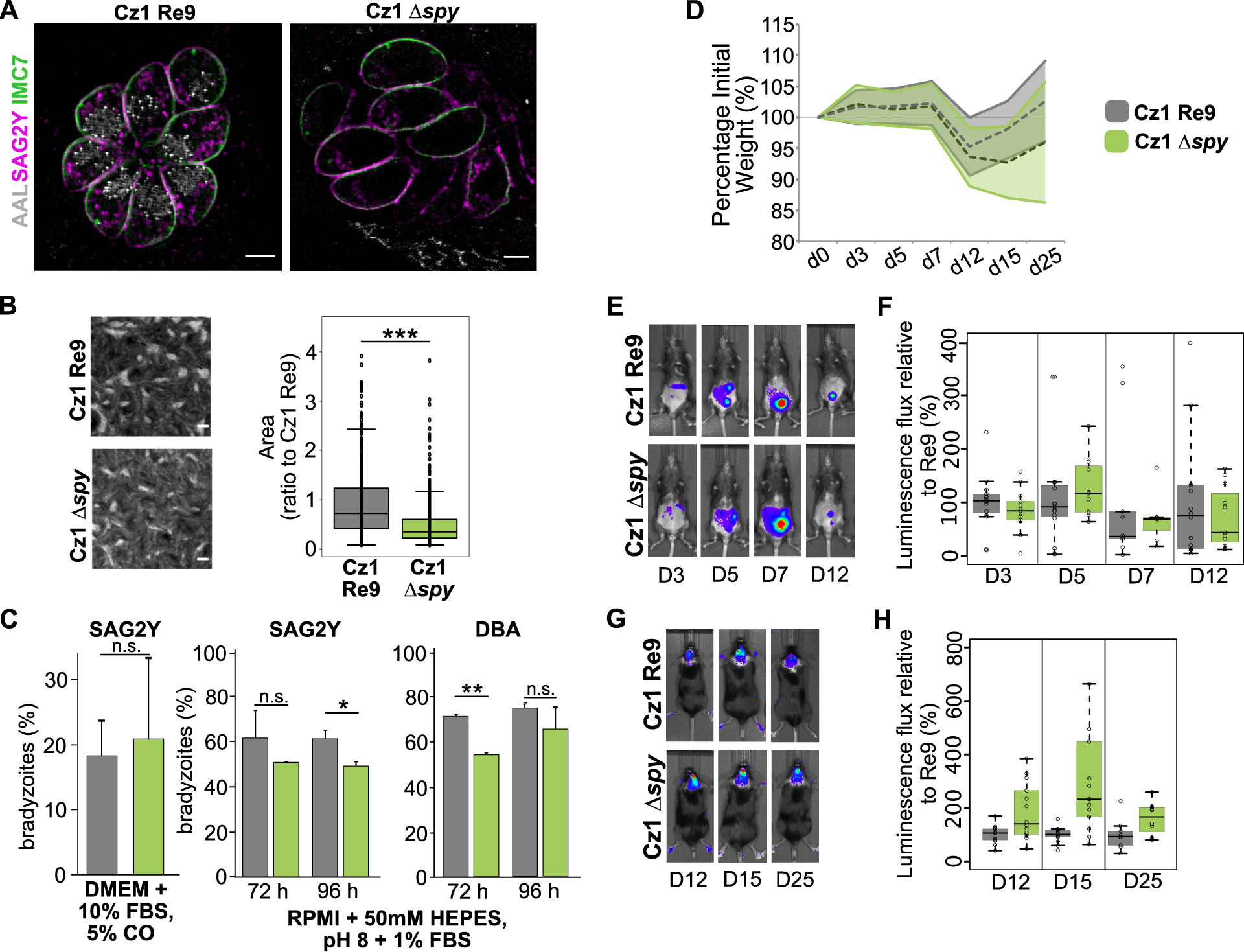
Analysis of *spy* knockout in *Toxoplasma* type II strain CZ1 *in vitro* and *in vivo*. *A*. The CZ1 Re9 (parental strain) and the derivative CZ1 Δ*spy* strains were immunostained with SAG2Y, which confirmed their differentiation to bradyzoites. AAL indicates *O*-Fuc status, while IMC7 outlines the cells owing to its localization to the inner membrane complex near the plasma membrane. Scale bar: 5 μm. *B*. Clonal plaque assays indicated a stronger growth defect in Δ*spy* in type II CZ1 than in type I RH (*** *p* value = 2×10^-16^) (Fig. 3F). Three biological repeats. Scale bar: 0.2 cm. *C*. Spontaneous differentiation (left) from tachyzoites to bradyzoites in DMEM + 10% FBS was not affected by Δ*spy* as determined by SAG2Y labeling. In contrast, differentiation into bradyzoites by incubation in alkaline induction media (pH 8 and 1%FBS) lowered the percentage of bradyzoites at 72 h (measured by DBA labeling) or 96 h (based on SAG2Y antibody) (* *p* < 0.01; ** *p* <0.005). Quantification performed on 100 vacuoles/cell line/experiment, ± standard deviation; three biological repeats). *D*. Parental strain (CZ1 Re9) and Δ*spy* each lost weight after intraperitoneal injection of 10^4^ tachyzoites (sublethal infection). *E*. Luminescence in the abdominal cavity (acute phase) was measured at days 3, 5, 7 and 12. Representative examples are shown *F*. Summary of results from three biological repeats, with 5 mice per repeat, showed no difference between parental strain and Δ*spy. G*. Representative examples of luminescence measured during brain infection on days 12, 15 and 25. *H*. Summary of results from three biological repeats, with 5 mice per repeat, showed increased luminescence in brains of the Δ*spy* compared to parental strain.

When cloned on fibroblast monolayers, plaques formed by Δ*spy* CZ1 were on average 50% smaller than those of the parental Re9 strain (Fig. 5B), pointing to a stronger growth defect in this type II strain than observed for the type I RH strain (Fig. 2F). Using SAG2Y as a bradyzoite marker, we found no difference in spontaneous differentiation of Δ*spy* and Re9 strains (Fig. 5C). However, when parasites were cultured in alkaline medium to induce differentiation and analyzed 72 or 96 h post incubation, a modest reduction in the number of bradyzoites was observed for Δ*spy*, based on either SAG2Y or DBA labelling (Fig. 5C).

C57BL/6 mice infected by intraperitoneal injection with 10^4^ tachyzoites from Re9 parental and Δ*spy* CZ1 strains showed similar patterns of weight loss and recovery on days 3, 5, 7, 12, 15 and 25 post infection (Fig. 5D). While there was no statistical difference in the luminescence signal from the abdominal cavity during the acute phase of infection (Figs. 5E and 5F), Δ*spy* parasites displayed consistently higher signals than the parental CZ1 Re9 strain from the brain during infection of the brain (Figs. 5G and 5H). As these were sublethal doses of infection, there was no difference in mortality between Re9 and Δ*spy* parasites (data not shown).

In summary, deletion of *Tg*SPY in type II CZ1 strain caused modest decreases in growth and differentiation in culture and a moderate increase in luminescence of parasites in mouse brains.

## Discussion

SPY is conserved in plants and algae, where it contributes to signaling and growth (26, 27). This report shows that SPY also occurs and plays key roles in a wide range of other organisms. Arguing for an important role for the *Tg*SPY OFT is the large number of proteins with SRDs modified by *O*-Fuc, many of which are predicted to be involved in gene transcription, protein transport, and signaling (22). Indeed, in a CRISPR/Cas9 screen examining growth of RH strain tachyzoites in culture, 69 AAL-enriched proteins, which include 33 with *O*-Fuc peptides identified directly by mass spectrometry, have average phenotype scores that are similar to those of essential proteins (Fig. S3) (41). Even so, *Tg*SPY is not essential. This is shown by the viability of the Δ*spy* in both type I RH and type II CZ1 strains and is consistent with the fitness score of −0.11 (41). The conclusion that the *O*-Fuc modification is dispensable for the function and/or stability of its nucleocytosolic protein substrates is in contrast to two molecular observations. First, that addition of *O*-Fuc to the SRDs appears to direct proteins to assemblies within the nucleus and, secondly, loss of an SRD from GPN or the absence of *O*-Fuc on Nup68-YFP (an SRD-YFP reporter-construct), and possibly even on SPY itself, markedly decreases protein abundance.

Knockout of *Tg*SPY slows growth and differentiation of type II CZ1 strain in culture and increases the luminescence from parasites in mouse brains. We have not yet determined the effect of *Tg*SPY knockout on cat infections, but we have observed that AAL-labeled assemblies are absent in the periphery of meiotic nuclei of sporulating oocysts (22). Finally, addition of *O*-Fuc is the second cytosolic glycosylation system present in *T. gondii*, which also adds a pentasaccharide to a hydroxyproline residue on Skp1, an E3 ubiquitin ligase subunit (33).

*Tg*SPY and *At*SPY share a GT41 catalytic domain with *Hs*OGT, have many of the same residues important for catalytic function, but have 11 rather than 13.5 TPRs (26–28). Our mutagenesis studies show that the conserved OGT/OFT residues in the catalytic domain are also important for *Tg*SPY function (26, 34, 35). However, two of the positions targeted for mutagenesis (D619 and E692) are not strictly conserved in all OGT/OFT sequences (27), suggesting the possibility of intrinsic tolerance to amino acid changes. The residual activity observed *in vivo* for several of the point mutants is consistent with other studies that have noted greater activity of mutants in cell systems (*e*.*g*. (42)), likely because *in vitro* assays cannot comprehensively reproduce the conditions found in the cell environment (43). Additionally, while *At*SPY-3TPRs was active *in vitro* (26), we could not detect any AAL labelling in our *T. gondii* tachyzoites assay. This could be at least partially due to the difference in substrates: a peptide from *Arabidopsis* DELLA protein was used in the *At*SPY *in vitro* assay (26), while *Tg*SPY in the cell assay is acting on native proteins. As is the case with *Tg*SPY, *Hs*OGT modifies proteins with IDRs including FG-Nups, and many enzymes involved with gene transcription and mRNA processing (24, 25, 44). While DELLA and PPR5 (a core circadian clock component) have been shown to be modified by *At*SPY, the vast majority of nucleocytosolic proteins with *O*-Fuc of plants are uncharacterized, as secreted proteins, which have N-glycans abundantly decorated with fucose, dominate AAL pulldowns (26, 45).

While *Hs*OGT uses UDP-GlcNAc, which is responsive to the metabolic state of the cell (24), *Tg*SPY and *At*SPY use GDP-fucose, whose potential dependence on nutritional status is unclear. Although addition of *O*-GlcNAc is a dynamic modification, because of the reciprocal activity of *O*-GlcNAcase, we have no evidence for removal of *O*-Fuc in *T. gondii* (46). Finally, there is extensive crosstalk between *O*-GlcNAcylation and phosphorylation in host proteins. About 80 % of AAL-enriched proteins in *T. gondii* have also been reported to be phosphorylated (47, 48). Whether this overlap translates to functional crosstalk between *O*-Fuc and phosphorylation in the parasite remains to be determined.

The bright sub-nuclear envelope punctae labeled by AAL may for the most part be composed of FG-Nups, which are heavily modified with *O*-Fuc (22) and correlates with their *O*-GlcNAcylation in mammalian cells. However, the punctae, though near to the NPCs, are not in perfect register, and *O*-fucosylation also modifies many different proteins not thought to be associated with the nuclear pores. The nuclear assemblies of *O*-fucosylated proteins in *T. gondii*, whose *O*-fucosylation tends to occur along portions of their sequences predicted to be intrinsically disordered regions (IDRs), are morphologically reminiscent of assemblies of proteins with IDRs in the host nucleus (such as Cajal, PML, and histone locus bodies). However, there is no evidence for *O*-GlcNAcylation of biomolecular condensates or liquid-liquid phase separations (22, 49). Although this might be a failure of detection in part due to the absence of lectins and antibodies that bind *O*-GlcNAc as selectively as AAL binds *O*-Fuc, unique physicochemical properties of Fuc may predispose *O*-fucosylated IDRs to phase separate. Alternatively, it is possible that *T. gondii* proteins that recognize *O*-Fuc may be involved in forming nuclear assemblies, even though there is no evidence for host proteins that recognize *O*-GlcNAc in a comparable manner to how SH2 domains recognize phosphotyrosine, Finally, the non-NPC *O*-fucosylated proteins might be associated with the low level of AAL binding observed throughout the nucleoplasm.

In conclusion, the cell and molecular biology *Tg*SPY is complex, as profound effects of the OFT on reporter protein stability are not matched by modest effects on growth in fibroblasts or mice. This raises the possibility that *O*-fucosylation of SRDs and other sites results in reversible protein accumulation in puncta that serve as reservoirs near nuclear pores, where they would be strategically situated to facilitate export to the cytoplasm or association with incoming cargo as needed. Thus *O*-fucosylation might offer a mechanism for generating a reserve of critical proteins that offer resistance to stress and may not be detected in standard culture conditions or mouse infection models. Because of numerous methodologic advantages (*e*.*g*. the use of AAL to localize and purify proteins modified with *O*-Fuc and expression of full-length *Tg*SPY in bacteria or in *T. gondii* Δ*spy*), future studies of the *Tg*SPY OFT should provide insights into protein assembly, transcription, signal transduction, and pathogenesis in *T. gondii*. These studies may also suggest functions for *O*-Fuc in plants, which are more difficult to study using cell biological and biochemical methods, and may also suggest new functions for host OGT, which is an essential enzyme in mammalian cells.

## Experimental procedures

### Ethical Statement

Culturing and genetic manipulation of *Toxoplasma gondii* RH and CZ1 strains was approved by the Boston University Institutional Biosafety Committee. Mouse infections with *T. gondii* were approved by the Boston University Institutional Animal Care and Use Committee.

### *T*. *gondii* cell culture and manipulation

*T. gondii* RH and CZ1 strains were propagated by passaging in HFF cultured at 37°C, 5% CO_2_ in Dulbecco’s Modified Eagle Medium (DMEM) supplemented with 10% fetal bovine serum (FBS), GlutaMax and 100 units/ml penicillin and 100 μg/ml streptomycin (Gibco). Electroporation of tachyzoites was performed as previously described (22) using 10^6^ parasites per reaction. Bradyzoites were allowed to spontaneously differentiate under normal culture conditions (DMEM + 10%FBS, 5% CO_2_), or after culture under alkaline conditions (RPMI + 50 mM HEPES, pH 8 + 1 % FBS), and were observed at 72 h and 96 h by probing with anti-SAG2Y antibody or DBA lectin (see below).

### Generation of reporter and complemented cell lines

Generation of wild type RH strains expressing YFP-tagged Nup67 and Nup68, NLS-YFP, SRD-YFP and *Tg*GPN was previously described (22). All proteins were expressed under the tubulin promoter with the exception of *Tg*GPN and *Tg*GPNΔSRD where the endogenous 5’ UTRs were used. The same constructs and protocol were used to generate RH Δ*spy* parasites expressing Nup68-YFP and NLS-YFP.

*Tg*GPNΔSRD open-reading frame (ORF) was generated by gene synthesis (GenScript) and cloned into p*gpn*GoI3xMYC_3_/sagCAT using BglII and AvrII restriction sites. The plasmid is designed to encode a C-terminal extension beyond the native C-terminus: PR[MYC-tag]NG[MYC-tag]NGARAE[MYC-Tag] (50). The presence of the transgene was verified by PCR using primers P233 and P8. Parasites were electroporated with 20 μg DNA, selected with chloramphenicol and cloned by limiting dilution.

To remove the LoxP-flanked DHFR cassette, RH Δ*spy* parasites (31) were electroporated with p5RT70DiCre-DHFR, (a kind gift of Markus Meissner, Ludwig-Maximilians-Universität München) (51), recovered in a T25, and cloned by limiting dilution. Clones were randomly selected and tested for loss of resistance to pyrimethamine. The genotypes of the pyrimethamine-sensitive clones were verified by PCR using primers P206, P207, and P208.

The *E. coli* codon-optimized ORF of *Toxoplasma* SPY (Fig. S1) was cloned into ptubGoIMYC_3_/sagCAT (50) using BglII and AvrII restriction sites. The resulting plasmid (ptub*Tg*SPYMYC_3_/sagCAT) was used to complement RH Δ*spy*. Parasites were electroporated, selected and cloned as above. The presence of the ectopic copy of *Tg*SPY was verified by PCR using primers P9 and P198. The same protocol was used to complement RH Δ*spy* parasites with *Tg*SPY point mutant and truncated constructs. Additionally, the presence of expected point mutants in the complemented cell lines was verified by amplifying the ORF fragment by PCR from gDNA followed by Sanger sequencing. Primers sequences are listed in Table S1.

### CRISPR/Cas9-mediated disruption of *spy* in CZ1

The approach to generate the *spy* knockout is described in Figure S3C. Based on Me49 genome sequence, there is a SNP four bases upstream of the C-terminal PAM site for disrupting *spy* suggesting that the RH gRNA directing to this site might not work on a Me49-like sequence (31). For this reason, we designed an additional pU6 plasmid expressing an additional gRNA to target the Cz1 *spy* C-terminus. The protospacer was cloned using oligonucleotides P127-128 (spy_t2). Oligos were annealed and phosphorylated, and the resulting dsDNA was cloned in the BsaI-digested pU6Universal plasmid (38) to generate the plasmid pU6_spyt. To replace the gene of interest, the gra1 5’UTRs-LoxP-HXGPRT-LoxP-mGFP-gra2 3’UTRs expressing cassette was amplified by PCR using primers containing about 30-40 bp homology sequence to the double stranded break sites (P129 and P130, Table S1). The cassette was derived from (52) as described in Agop-Nersesian *et al*. (manuscript in preparation). Following PCR amplification, the fragment was cloned into pCR2.1TOPO (Invitrogen), generating pSPYKO_HXmGFP. A point mutation on the mGFP resulted in non-fluorescent parasites (mGFP*). The cassette also contains StuI and SfoI restriction sites to allow easy isolation of the linear DNA after plasmid amplification by restriction digestion and gel purification. About 20 μg each of pU6_spyt and pDG_Spy and at least 5× molar excess of purified recombination cassette were used to electroporate CZ1 Δ*hpt* Δ*ku80* Re9. Following selection with xanthine and mycophenolic acid, single clones were obtained by limiting dilution, and editing of the correct locus was verified by Southern blotting (see below).

### Site-directed mutagenesis and truncated constructs

A Q5® site-directed mutagenesis kit (New England Biolabs, NEB) was used to generate the *Tg*SPY point mutants and truncation constructs. Mutagenesis was performed directly on ptub*Tg*SPY3xMYC/sagCAT using primers P173/P174 (K793A), P175/P176 (H623A). P177/P178 (D619A), P226/P227 (E692K), P228/P229 (G695D), P179/P180 (3TPRs-*Tg*SPY), and P181/P182 (GT41 only) (Table S1). All constructs were verified by Sanger sequencing.

The same protocol was used for site-directed mutagenesis of pET-TEV_*Tgspy*, using primers that included silent restriction sites as an aid in screening (Table S1).

### Semiquantitative RT-PCR

About 10^7^ tachyzoites from RH parental, *Tg*GPN, or *Tg*GPNΔSRD-expressing strains were harvested and lysed in TriReagent (Invitrogen). RNA was extracted according to the manufacturer’s instructions and further cleaned with PureLink Mini RNA and the concentration was calculated on a Nanodrop (Thermo). Different amounts of total RNA (4, 16, 32, or 40 ng) were used for first strand synthesis using SmartScribe RT (Clontech) and Oligo(dT)23 VN primer (NEB). A minus RT control was also performed using the highest total RNA amount (40 ng). Primers (P233-P8) were used to amplify a 500-bp gene-specific fragment from cDNA, common to both the full length and truncated *Tg*GPN-MYC_3_ transgene, using OneTaq polymerase (NEB). RH cDNA was used as control for primer specificity, and the *T. gondii* GDP-mannose 4,6-dehydratase (*Tg*GMD) (P101-P102) served as positive control for cDNA synthesis. All reactions were analyzed by gel electrophoresis. Primers are listed in Table S1.

### Southern blot

Tachyzoites from CZ1 parental strain and Δ*spy* clones were lysed in DNAzol (Invitrogen) and genomic DNA was isolated according to the manufacturer’s instructions. Samples were treated with 0.1 mg/ml RNaseA (Sigma Aldrich) and genomic DNA quality was assessed by gel electrophoresis. gDNA (5 μg/lane) was digested with EcoRV, separated on a 1% agarose gel in Tris-acetate-EDTA buffer and blotted on a nylon membrane. A probe for mGFP ORF was generated using the PCR DIG Synthesis Probe Kit (Roche) and primers P35-P36 (Table S1) and the blot was processed and developed according to the manufacturer’s instructions.

### *Tg*SPY recombinant expression in *E*. *coli*

The predicted ORF for full-length *Tg*SPY from TGGT1_273500, annotated as an *O*-linked *N*-acetylglucosamine transferase in ToxoDB v46, was used to generate a cDNA that was codon-optimized (see Fig. S1) and synthesized by GenScript (Piscataway, NJ) for expression in *E. coli*. The *spy* cDNA was cloned into pET15-TEV, using the indicated restriction enzymes (Fig. S1), resulting in a version of *Tg*SPY that was N-terminally tagged with a His_6_ sequence followed by a TEV-protease site, fused to the authentic N-terminus of *Tg*SPY, including its Met. pET-TEV_*Tgspy* was electroporated into *E. coli* Gold and its expression was induced in the presence of 0.5 mM IPTG for 18 h at 22 °C. Cells were collected and protein was extracted and purified over a Ni^+2^-column and a Superdex 200 column, essentially as described previously for His_6_Skp1 (53). A highly enriched sample from an included volume fraction was analyzed.

For improved expression of the mutant proteins, an autoinduction method was used following electroporation into *E. coli* (54). After 42 h of growth at 18 °C (no IPTG induction), extracts were prepared as above, and the SPY protein was captured on a Co^++^-Talon column. After washing the columns with 30 mM imidazole in 50 mM HEPES-NaOH (pH 7.4), 0.5 M NaCl, the SPY fraction was eluted with 250 mM imidazole in the same buffer, immediately dialyzed at 4° C against 50 mM HEPES-NaOH (pH 7.4), concentrated by centrifugal ultrafiltration, and frozen at −80 °C in aliquots. Samples were analyzed by SDS-PAGE and staining with Coomassie blue, or Western blotted and p probed with anti-His Ab, as described (33).

### GST-SRD expression in *E*.*coli*

The sequence encoding for the 79 aa at the N-terminus of *Tg*GPN (TGGT1_285720) was used to generate a cDNA that was codon-optimized for *E. coli* and synthesized by GenScript to include BamHI and XhoI restriction sites. The SRD cDNA was cloned via BamHI/XhoI into pGEX6P-1, resulting in a N-terminal GST tag followed by the PreScission protease cleavage site and the SRD encoding sequence. The final pGEX6P-1-SRD plasmid was used to transform chemically competent *E. coli* BL21(DE3) and expression was induced with 0.25 mM IPTG for 3.5 h at 37°C. Cells were collected and lysed by sonication in the presence of lysozyme. The soluble fraction was obtained by centrifugation and the recombinant protein was purified by anion exchange chromatography on a Q-Sepharose column.

### Recombinant *Tg*SPY activity assay

Purified His_6_*Tg*SPY was assayed for sugar nucleotide hydrolysis activity and transferase activity for transfer of Fuc from GDP-Fuc to protein acceptors. The Standard Enzyme Buffer (SEB) consisted of 50 mM 2-(*N*-Morpholino)ethanesulfonic acid (MES) pH 6.5, 50-70 mM NaCl, 5 mM MgCl_2_, 2 mM DTT, 10 µg/ml aprotinin, 10 µg/ml leupeptin, 1 mM PMSF. Hydrolysis of UDP-Gal, UDP-GlcNAc, GDP-Fuc and GDP-Man was quantified using UDP-Glo and GDP-Glo assays (Promega), as described (32).

Transferase activity was assayed as transfer of Fuc from GDP-Fuc to protein. Typical assays contained 2 µM GDP-[^3^H]Fuc (22,400 dpm/pmol; American Radiochemical Corporation) in SEB and were incubated for 0 or 7 h at 29°C. Auto-fucosylation reactions were conducted for 7 h in the presence of purified His_6_*Tg*SPY and unlabeled GDP-Fuc. Fucosylation was assayed by SDS-PAGE and Western blotting with biotinylated AAL as described (22). Scout assays indicated that activity was highest at pH 6.5, insensitive to NaCl from 20 to 150 mM, not affected by addition of 2 mM MnCl_2_, and not significantly affected by 0.2% Tween-20 or 2 mg/ml BSA. To assay transfer to a second protein, the candidate acceptor protein GST-SRD was expressed in *E. coli* and purified using Q-Sepharose chromatography as described above. The reaction was conducted in the presence of 2 µM GDP-[^3^H]Fuc (22,400 dpm/pmol; American Radiochemical Corporation), and incorporation was assayed by resolving the reaction products by SDS-PAGE and counting gel slices for radioactivity in a liquid scintillation counter, as described (55). To test incorporation into native *Toxoplasma* proteins, desalted cytosolic extracts of tachyzoites were prepared as before (55) and used to assay incorporation into native acceptor substrates, using the same SDS-PAGE assay.

To assay transfer to a peptide, the tryptic peptide ZN190 (SSSSSASSSSSSFPSSSSSDSVPPR), which derives from *Tg*RINGF1 (TGGT1_285190) and has been reported to be *O*-fucosylated in cells (22), was synthesized (GenScript) and dissolved at 10 mM in 50 mM NH_4_HCO_3_. Typical reactions contained, in 20 µl, 0.1 mM peptide, 10 µM GDP-Fuc (0.1 µCi GDP-[^3^H]Fuc), 50 mM MES-NaOH (pH 6.5), 50 mM NaCl, 3 mM NaF, 2 mM DTT, 1 mg/ml bovine serum albumin, 10 µg/ml aprotinin, 10 µg/ml leupeptin, and were incubated for 40 min at 29 °C. Reactions were stopped by the addition of 500 µl of 50 mM formic acid, 1 M NaCl, and stored at –20 °C. Samples were applied under vacuum to 50 mg C_18_ SepPak cartridges (Waters WAT054955) that had been precycled with 6 × 1 ml MeOH, 2 × 1 ml 50 mM formic acid, and 4 × 1 ml 50 mM formic acid, 1 M NaCl. The sample tubes were rinsed with 1.5 ml 50 mM formic acid, 1 M NaCl and the mixture was applied to the cartridge, which was then washed with 12 × 1 ml 50 mM formic acid, 1 M NaCl, and 2 × 1 ml 50 mM formic acid. The final two washes were collected in a 20-ml scintillation vial and mixed with 15 ml of BioSafe-II (RPI Corporation, 111195), and referred to as ‘W’. The cartridge was eluted with 2 × 1 ml MeOH into a 20-ml scintillation vial and mixed with 15 ml BioSafe-II, referred to as E1. A second elution with MeOH yielded negligible signal (dpm). All samples were assayed for radioactivity in a Beckman LS6500 liquid scintillation counter. Data was initially recorded as ‘W-E’ dpm. Values for ‘W’ typically did not exceed 3% of ‘E’ values for positive reactions with the ZN190 peptide. Matched time zero values, which typically did not exceed 10% of final values and were similar in reactions lacking added peptide or *Tg*SPY, were subtracted from the timed reaction values. Incorporation was enzyme concentration and time dependent under the conditions used (data not shown).

### Reaction product characterization

Autofucosylation of His_6_*Tg*SPY was performed in the presence of GDP-[^3^H]Fuc, as described above for GST-SRD in the presence of 0.1U/μl antarctic alkaline phosphatase, and the reaction was subjected to SDS-PAGE and electroblotted onto a PVDF membrane as described (67). The membrane position corresponding to SPY was excised with a scalpel and subjected to conditions of reductive β-elimination as reported previously (31). Briefly, the membrane fragment was rinsed with H_2_O, suspended in 200 µl of 0.05 M NaOH, 1 M NaBH_4_, and incubated at 45 °C for 16 h. Unreacted NaBH_4_ was neutralized by the addition of 10% acetic acid, and Na^+^ was removed by passage over a 1-ml Dowex 50W-XB (H^+^ form, Sigma-Aldrich) column equilibrated with 5% acetic acid. The eluate was dried under a N_2_ stream at 45 °C. The residue was dissolved in 750 µl of a 9:1 mixture of methanol:acetic acid and dried under vacuum centrifugation, repeated two times, to remove boric acid. Three successive additions of 750 µl H_2_O followed by vacuum centrifugation were conducted before analysis on a Thermo Scientific ICS 5000+ High Performance Anion Exchange Chromatography (HPAEC) system. A mixture of 2.5 nmol myoinositol, and 5 nmol each of fucitol, sorbitol and mannitol, was added to the sample, which was injected onto a CarboPac MA-1 (Dionex) column and eluted under isocratic conditions with 612 mM NaOH at 0.4 ml/min, with pulsed amperometric detection. Fractions were collected manually, neutralized with an equal volume of 612 mM acetic acid, mixed with 7 ml Biosafe II scintillation fluid (Research Products International), and counted in a liquid scintillation counter (Beckman LS 6500).

### Immunofluorescence analysis

Intracellular tachyzoites were fixed in 4% paraformaldehyde (PFA) in phosphate-buffered saline (PBS) for 20 min at room temperature (RT), permeabilized in 0.25% TX-100 in PBS for 15 min at RT and blocked in 3% bovine serum albumin (BSA) in PBS for 1 h, at RT. RH Δ*spy* tachyzoites were transiently electroporated with ptubSRDYFP/sagCAT and fixed 24 h post electroporation.

Primary antibodies were used at the following concentrations: mouse anti-c-MYC 9E10 (DHSB) 4 μg/ml, mouse anti-epichromatin PL2-6 1:100 (56), rabbit anti-β-tubulin 1:1000, rat anti-IMC7 1:1000, and rabbit anti-SAG2Y 1:5000. The antibodies against epichromatin, IMC7, and β-tubulin were a kind gift of Marc-Jan Gubbels (Boston College), while the anti-SAG2Y was a kind gift of Jeroen Saeij (University of California Davis). Secondary goat AlexaFluor-conjugated anti-mouse, anti-rat, and anti-rabbit antibodies (Molecular Probes) were used at a 1:800 dilution. *Aleuria aurantia* lectin (AAL) and *Dolichos biflorus* agglutinin (DBA), purchased from Vector Labs, were conjugated to AlexaFluor −594 or −488 succinimidyl esters (Molecular Probes) following the manufacturer’s instructions and used at a 1:250 and 1:125 dilution, respectively. Finally, nuclei were labeled with 1 μg/ml 4’,6-diamidino-2-phenylindole (DAPI) for 10 min at RT. Coverslips were mounted using Vectashield (Vector Labs) mounting medium.

For counting the number of vacuoles positive for SAG2Y or DBA, cells were observed at a total magnification of 400x on a Zeiss AXIO inverted microscope with a Colibri LED operated via ZEN software. A minimum of 100 vacuoles/biological repeat were counted for each strain at each time point and the average ± SD of three independent repeats is shown. A two-tailed, homoscedastic Student t test was used to calculate the *p* values.

Super-resolution structured illumination microscopy (SIM) was performed on a Zeiss ELYRA microscope. Images were acquired with a 63x/1.4 oil immersion objective, 0.089 µm z sections at RT and processed for SIM using ZEN software. For single optical sections, images were further processed with Fiji (57).

### Quantification of AAL staining and AAL-positive puncta

To quantify and compare the intensity of AAL staining between wild type and complemented parasites, images were taken at the ELYRA microscope as detailed above and processed by SIM. A maximum intensity projection was then generated using the ZEN software. In Fiji, the epichromatin staining was used to define the nuclear region (NRoI) and the mean intensity of the NRoI in the AAL channel was measured. The ratio to wild type staining from an average of three biological repeats is shown. A total of 111 tachyzoites from the parental strain and 106 from Δ*spy*::*Tg*SPY were quantitated.

The number of AAL-positive puncta was counted using the ‘Analyze Particle’ function in Fiji (57). The puncta from a total of 86 tachyzoites from the parental strain and 104 from Δ*spy*::*Tg*SPY were quantitated. Box plots and statistical analyses (Student t Test) were performed using R Studio.

### *T*.*gondii* tachyzoites total cell lysate

Total cell lysates were extracted from extracellular tachyzoites. Parasites were harvested by centrifugation, washed twice in PBS and lysed in 1x reducing SDS-PAGE loading buffer containing 0.1M DTT. Lysates were heated 10 min at 96 °C. For AAL blotting of tachyzoites, the protocol was modified as follows: parasites were washed four times in PBS before lysis and heating was performed at 50 °C for 20 min.

### Western and lectin blot analysis

About 5×10^6^ cells equivalent/lane *T. gondii* tachyzoites were loaded on 8-16% TGX gels (Life Technologies). After SDS-PAGE separation, proteins were blotted on PVDF and then the membranes were blocked in 50 mM TrisHCl, 0.15 M NaCl, 0.25% BSA, 0.05% NP-40 pH 7.4. β-elimination on blot was performed by incubating the membrane 16 h in 55 mM NaOH at 40 °C, under rotation, before blocking as described above (58). Both primary and secondary antibodies were diluted in blocking buffer as follows: mouse MAb anti-cMYC 9E10 0.4 μg/ml, mouse MAb anti-α-tubulin 12G10 1:1000 (DHSB), biotinylated-AAL (Vector Labs) 2 μg/ml, mouse anti-GFP (Roche) 1:2000, ExtrAvidin-HRP (Sigma Aldrich) 1:10000 and goat anti-mouse HRP-conjugated (BioRad) 1:1000. For AAL inhibition, the biotinylated lectin at its working dilution was incubated 30 min at RT with 0.2 M methyl-α-L-fucopyranoside prior to blot incubation. Blots were developed for detection by chemiluminescence (SuperSignal West Pico PLUS) using an ImageQuant LAS4000 imager (GE Healthcare) and quantification was performed using the ImageQuant TL software. The average of three biological repeats ± SD is shown.

### Flow cytometry

Between 5×10^7^-1×10^8^ *T. gondii* extracellular tachyzoites expressing either full length or truncated *Tg*GPN were harvested by centrifugation, washed twice in PBS and fixed in 4% paraformaldehyde (PFA) in PBS for 30 min at room temperature (RT). From this point forward, all wash steps were performed by centrifugation for 5 min at 500 x *g*. Parasites were permeabilized in 0.1% TX-100 in PBS for 10 min, with rotation at RT and blocked in 3% BSA in PBS for 1 h, rotating at RT. Each sample was divided into three aliquots and these were incubated with either AlexaFluor488-conjugated AAL (1:250), mouse anti-c-MYC 9E10 (4 μg/ml) followed by goat anti-mouse AlexaFluor488 (1:500) or buffer alone (3%BSA in PBS) as unlabeled control. All incubations were performed at RT, rotating in the dark for 1 hr. Cells were resuspended in PBS and analyzed on a FACSCalibur (BD).

### Plaque assay

Host cell monolayers on 6-well plates were infected with 250 parasites/well (RH) or 1000 parasites/well (CZ1) of either parental, Δ*spy* or complemented strains. RH and CZ1 parasites were allowed to grow at 37 °C, 5% CO_2_ for 5 or 8 days, respectively. After extensive washing in PBS, cells were fixed in ice-cold methanol at −20 °C for at least 20 min and stained with 2% crystal violet for 15 min. Wells were then washed with PBS, air-dried and imaged on an ImageQuant LAS400. Plaque areas were measured with Fiji and the box plot shows the average of three biological repeats ± SD. A two-tailed, homoscedastic Student t test was used to calculate the *p* values (R Studio).

### *In vitro* differentiation

HHF were grown to confluence on glass coverslips in 12-well plates. CZ1 parental and Δ*spy* strains were inoculated with a multiplicity of infection (MOI) of 0.5. When differentiation was evaluated under normal culture conditions, parasites were allowed to replicate for 72 h before fixation. Alternatively, parasites were allowed to replicate for 24 h before exchanging to alkaline stress medium: RPMI 1640 (Gibco) supplemented with 1% FBS, 100 units/ml penicillin, and 100 μg/ml streptomycin and buffered with 50 mM HEPES (pH 8.1). Parasites were incubated at 37 °C in the absence of CO_2_ for 72 or 96 h before fixation. Fixation and labelling were performed as described above.

### Mouse infections

Female C57BL/6 5 week-old mice were purchased from Charles River Laboratories (Wilmington, MA) and housed 5/cage with *ad libitum* access to food and water. Infections were performed after one week of acclimation as described (Agop-Nersesian *et al*., manuscript in preparation). Briefly, CZ1 parental and Δ*spy* strains were harvested immediately before infection from one T25 by scraping HFF monolayers. Tachyzoites were released from PSVs by passing twice through a 27G needle attached to a 5-ml syringe. Host cell debris was removed by filtering through a 3-μm polycarbonate membrane (Millipore). Parasites were counted using a haemocytometer and diluted to be 5×10^4^/ml. Each mouse was injected intraperitoneally (I.P.) with 200 μl of the suspension (10^4^ tachyzoites). Additionally, for each experiment a plaque control was run: the same parasite suspension was used to infect the 6-well plate with 1000 and 2000 parasites of parental and knockout strains. Plaques were allowed to form and monolayers were fixed, labelled and quantitated as above. Mice were weighed on the day of infection and of IVIS measurements. All mice were sacrificed 5 weeks post infections.

### *In Vivo* Imaging

*In vivo* luminescence measurements were performed as described in Agop-Nersesian *et al*., (manuscript in preparation). C57BL/6 black mice were shaved the day before measurement on day 3 (abdomen) and day 12 (back of the head). If necessary, additional shaving was performed at later time points. Measurements were taken on days 3, 5, 7, 12, 15 and 25 post infection. A sterile solution of D-luciferin (PerkinElmer, Waltham MA) in Mg^2+^- and Ca^2+^-free DPBS was injected I.P. at 300 mg/kg, Mice were then sedated in a XGI-8 Anaesthesia System with 1.5% isofluorane and moved to the chamber of a IVIS Spectrum *In vivo* Imaging System (PerkinElmer) under 0.25% isofluorane. Abdomens were imaged 10 min post luciferin injection with exposures between 30 sec and 2 min. For brain imaging, a second luciferin injection was administered 20 min after the first one and, following the same sedation procedure, mice were imaged 30 min after the second injection with a 5-min exposure. Images were processed and quantified on Living Image (PerkinElmer) as total flux (photons/seconds) and adjusted to the numbers from the plaque assays to account for discrepancies in the number of parasites used to infect the mice. Bioluminescence measurements are presented as flux relative to the parental strain.

## Acknowledgements

We are grateful to M. Osman Sheikh for conducting the NDP-sugar Glo assays and Angela Park for preparing GST-SRD. We thank Marc-Jan Gubbels (BC), Jeroen Saeij (UC Davis) and Markus Meissner (LMU Munich) for generously providing reagents. We thank Bret Judson and the Boston College Imaging Core for infrastructure and support (National Science Foundation Grant 1626072). Finally, a shout out to Phil Robbins, the best collaborator ever, with whom we initiated these studies.

## Author contributions

G.B., C.M.W., and J.S. designed the research; G.B., C. A-N., H.vdW., M.M., and H.K. performed the research; G.B., C.A-N., H. vdW., C.M.W., and J.S. analysed the data; G.B., C. A-N., C.M.W. and J.S. wrote the paper. All authors read and approved the finalized manuscript.

## Abbreviations

AAL: *Aleuria aurantia* lectin
*At*SPY: *Arabidopsis thaliana* SPINDLY
CAZy: Carbohydrate Active EnZyme
DBA: *Dolichos biflorus* agglutinin
SPINDLY; FG-Nup: Phe/Gly-repeats nucleoporin
GT41: glycosyltransferase 41
GST: glutathione-S-transferase
HFF: human foreskin fibroblast
IDR: intrinsically disorder region
Me-αFuc: α-methyl fucopyranoside
NLS: nuclear localization signal
NPC: nuclear pore complex
*O-*Fuc: *O-*linked fucose
OFT: *O*-fucosyltransferase
OGT: *O*-GlcNAc transferase
SIM: structured illumination microscopy
SRD: serine-rich domain
*Tg*GMD: *Toxoplasma gondii* GDP-mannose 4,6-dehydratase
*Tg*GPN: *Toxoplasma gondii* GPN-loop GTPase
*Tg*GPNΔSRD: *Tg*GPN missing the SRD at its N-terminus
*Tg*SPY: *Toxoplasma gondii* SPINDLY
*Tg*SPY-3TPRs: *Tg*SPY with three C-terminal TPRs
TPR: tetratricopeptide repeat
YFP: yellow fluorescent protein

