## Supporting Information for "The nucleocytosolic *O*-fucosyltransferase *Spindly* affects protein expression and virulence in *Toxoplasma gondii*"

This Supporting Information file includes:

**Table S1**

**Figures S1-S3**

**Table S1**. Primers used in this study

| **Primer** | **Number** | **Sequence** | **Restriction sites** | **Ref.** |
| --- | --- | --- | --- | --- |
| gmdF | P101 | CGGATCCAAAATGGAAGGCGAAAACG |  |  |
| gmdR | P102 | CCTAGGCGCCTTTTGGACTCTCC |  |  |
| SPYt2_F | P127 | AAGTTGAGCTGCTAGCTCGCAAACACG |  |  |
| SPYt2_R | P128 | AAAACGTGTTTGCGAGCTAGCAGCTCA |  |  |
| SPYKO_RO_F | P129 | AGGCCTCACGCCGGAGTCTCCTCCTCACAGTCGCGGCCGTTCtcgaaggctgctagtactg | StuI |  |
| SPYKO_RO_R | P130 | GGCGCCAGACTCTTGATCGCCTCTGCAGCGCTCTGGCTgcttaacaccattgcattcc | SfoI |  |
| TgSPY_F | P161 | GAATGGACCTGATCCTTCC |  |  |
| TgSPYe3_R | P162 | TGGTTGGTGAGGATTCG |  |  |
| K793A_F | P173 | CAACCTGGCGgcaCTGGGTAACC |  |  |
| K793A_R | P174 | TTGAAGCTACCGAAGGTAAC |  |  |
| H623A_F | P175 | CTTCTTCACCgcaAGCGTCAGCTACTTCATCC |  |  |
| H623A_R | P176 | TCCGGACCAATGTAGCCA |  |  |
| D619A_F | P177 | CATTGGTCCGgcaTTCTTCACCCACAG |  |  |
| D619A_R | P178 | TAGCCAACGCGGATCACA |  |  |
| 3T_F | P179 | GCGGAAGCGTACAACAAC |  |  |
| 3T_R | P180 | CATAGATCTAAAAGGGAATTCAAGAAAAAATG |  |  |
| gt41_F | P181 | AACGCGTTCCACAACAAAC |  |  |
| gt41_R | P182 | CATAGATCTAAAAGGGAATTCAAG |  |  |
| gt41_dia_R | P197 | GCAGGGTCTTGAACATCTG |  |  |
| tgmut_F | P198 | CCAGGGTGACATCGAGGACA |  |  |
| 273500_F2 | P206 | ATGAATGGACCTGATCCTTCCC |  | 31 |
| 273500_R | P207 | CTAGCAGCTGGCGATACAGC |  | 31 |
| DHFR_seqR | P208 | CGAATGACACACAGGAACTACGC |  | 31 |
| E692K_F | P226 | CATTCTGGTCAAGCTGGCAGGTC |  |  |
| E692K_R | P227 | TCGATACGGTCTTCCTCG |  |  |
| G695D_F | P228 | GAACTGGCAGaTCACACCGCA |  |  |
| G695D_R | P229 | GACCAGAATGTCGATACGGTC |  |  |
| 720F | P233 | GAGCAACATGCTGTACGCAT |  |  |
| GFP_f | P35 | AGCTCGAGATGAGTAAAGGAGAAGAAC | XhoI |  |
| GFP_r | P36 | GCTTAATTAACTATTTGTATAGTTCATCCATG | PacI | 21 |
| SRD_F | P7 | CTAAAATGGAGACCTCTTCTC |  | 22 |
| cMycR | P8 | CACCGTTCAAGTCTTCCTCGG |  | 22 |

| *E. coli* |
| --- |

| Tg-D619A-Q5S |  | GCTACATTGGgcccgcCTTCTTCACCCAC | ApaI |
| --- | --- | --- | --- |
| Tg-D619A-Q5AS |  | CAACGCGGATCACACGGC |  |
| Tg-H623A-Q5S1 |  | CTTCTTCACCgctAGCGTCAGCTACTTCATC | NheI |
| Tg-H623A-Q5AS |  | TCCGGACCAATGTAGCCA |  |
| Tg-E692K-Q5S |  | CATTCTGGTCaagcttGCAGGTCACAC | HindIII |
| Tg-E692K-Q5AS |  | TCGATACGGTCTTCCTCG |  |
| Tg-G695D-Q5S |  | cagaTCACACCGCACACAACCG | NheI(split) |
| Tg-G695D-Q5AS |  | ctagcTCGACCAGAATGTCGATACG | NheI(split) |
| Tg-K793A-Q5S |  | CAACCTGGCGgccctaGGTAACCAGG | AvrII |
| Tg-K793A-Q5AS |  | TTGAAGCTACCGAAGGTAAC |  |

**FIGURES**

**Figure S1. *Tg*SPY cDNA sequence.** Predicted native coding sequence, based on TGME49_273500, is in black; synthetic, which includes cDNA sequence and 3’ UTR, after cloning in pET15-TEV is in blue; calculated amino acid sequence of the synthetic cDNA is in bold black. The native start and stop codons are highlighted in teal green. Restriction sites are marked in khaki (Bgl*II*), blue (Nco*I*), grey (Nde*I*), yellow (BamH*I*), new color (Spe*I*), and green (Hind*III*).

```
                                             M  G  S  S  H
                                     gatataccatgggcagcagccat

H  H  H  H  H  S  S  G  R  E  N  L  Y  F  Q  G  H  M  R  S
catcatcatcatcacagcagcggcagagaaaacttgtatttccagggccaTATGAGATCT

M  N  G  P  D  P  S  R  P  C  L  T  P  E  S  P  P  H  S  R     20
ATGAATGGACCTGATCCTTCCCGGCCGTGCCTCACGCCGGAGTCTCCTCCTCACAGTCGC   60 Tgspy
ATGAACGGTCCGGACCCGTCTCGTCCGTGCCTGACCCCGGAGAGCCCGCCGCACAGCCGT      synthetic

G  R  S  C  P  P  S  D  Q  L  G  N  S  S  S  S  S  A  T  T     40
GGCCGTTCCTGCCCTCCCTCGGATCAACTTGGCAATTCCTCTTCGTCTTCTGCTACCACA  120
GGCCGCAGCTGCCCGCCGAGCGACCAGCTGGGTAACAGCAGCAGCAGCAGCGCAACCACC

P  A  T  D  L  D  S  A  V  Q  D  N  I  F  A  Q  A  P  S  E     60
CCGGCAACCGACTTGGACTCTGCAGTGCAGGACAACATCTTTGCGCAGGCGCCCTCGGAA  180
CCGGCAACCGACCTGGACAGCGCGGTGCAGGACAACATCTTCGCACAGGCACCGAGCGAA

T  S  S  R  P  L  P  I  P  A  A  N  A  L  R  H  L  P  S  P     80
ACTTCTTCTCGTCCTCTTCCGATCCCTGCAGCCAACGCCCTGCGACACCTGCCTTCGCCG  240
ACCAGCAGCCGTCCGCTGCCGATTCCGGCAGCGAACGCGCTGCGTCACCTGCCGAGCCCG

P  L  A  R  E  A  C  D  G  G  A  R  P  S  P  Q  R  N  C  C    100
CCGCTCGCCAGGGAAGCCTGTGACGGAGGCGCGAGGCCCTCGCCTCAGCGGAACTGCTGC  300
CCGCTGGCACGTGAGGCGTGCGACGGCGGTGCGCGTCCGAGCCCGCAGCGCAACTGCTGC

D  G  R  P  A  A  C  N  H  R  A  V  S  D  V  C  A  S  L  P    120
GACGGCAGGCCGGCTGCATGCAACCACCGTGCAGTCAGCGACGTGTGTGCGTCGCTCCCA  360
GACGGCCGTCCGGCGGCGTGCAACCACCGTGCAGTTAGCGACGTCTGCGCAAGCCTGCCG

E  K  S  F  A  R  H  F  L  T  S  S  G  T  F  P  S  A  A  E    140
GAAAAGTCCTTTGCTCGTCATTTCCTCACGTCTTCTGGGACCTTTCCGTCTGCTGCGGAG  420
GAAAAGAGCTTCGCGCGTCACTTCCTGACCAGCAGCGGTACCTTCCCGAGCGCAGCAGAG

I  L  K  K  A  A  F  F  N  S  G  N  R  P  H  D  A  L  L  L    160
ATCCTAAAGAAAGCGGCGTTCTTCAACTCGGGCAACCGCCCTCATGATGCCCTTCTCCTC  480
ATCCTGAAGAAAGCGGCGTTCTTCAACAGCGGTAACCGTCCGCACGACGCACTGCTGCTG

C  N  A  G  L  E  V  Y  A  E  D  A  D  L  W  N  C  K  G  V    180
TGCAACGCTGGCCTGGAAGTTTACGCCGAAGATGCCGACTTGTGGAACTGCAAAGGCGTC  540
TGCAACGCGGGTCTGGAAGTGTACGCGGAAGACGCGGACCTGTGGAACTGCAAAGGCGTG

T  L  R  A  L  G  R  L  Q  E  A  L  D  C  C  R  E  A  L  R    200
ACTCTTCGAGCTCTTGGAAGACTCCAGGAGGCACTTGATTGCTGCCGAGAGGCGCTTCGT  600
ACCCTGCGTGCGCTGGGTCGCCTGCAGGAAGCGCTGGACTGCTGCCGTGAGGCACTGCGT

L  D  P  G  N  T  N  A  L  N  N  I  G  V  A  L  K  E  R  G    220
CTTGATCCAGGGAATACAAATGCTCTAAACAACATTGGAGTTGCATTGAAGGAGAGAGGC  660
CTGGACCCGGGCAACACCAACGCGCTGAACAACATCGGCGTTGCGCTGAAGGAGCGTGGT

E  L  L  Q  A  V  E  H  Y  R  A  S  L  V  A  N  P  H  Q  P    240
GAACTTCTACAAGCTGTCGAGCACTACCGGGCCTCTCTGGTGGCGAATCCTCACCAACCA  720
GAACTGCTGCAGGCGGTCGAACACTACCGTGCAAGCCTGGTGGCGAACCCGCACCAGCCG

T  C  R  T  N  L  A  V  A  L  T  D  L  G  T  K  L  K  Q  E    260
ACATGCCGCACAAACTTGGCAGTTGCCCTCACCGACTTGGGCACGAAGCTGAAGCAGGAG  780
ACCTGCCGCACCAACCTGGCAGTTGCACTGACCGACCTGGGCACCAAGCTGAAACAGGAA

K  K  L  Q  A  A  L  V  C  Y  T  E  A  L  T  A  D  P  T  Y    280
AAGAAGCTGCAGGCAGCGCTCGTTTGCTACACAGAGGCACTGACCGCAGACCCGACCTAC  840
AAGAAACTGCAGGCAGCACTGGTGTGCTACACCGAAGCACTGACCGCGGACCCGACCTAC

A  P  C  Y  Y  N  L  G  V  I  H  A  E  T  D  D  P  H  T  A    300
GCGCCTTGCTACTACAACCTTGGCGTTATCCACGCAGAAACAGATGACCCCCACACCGCT  900
GCGCCGTGCTACTACAACCTGGGTGTGATTCACGCAGAGACCGACGACCCGCACACCGCG

L  Q  M  Y  R  E  A  T  R  L  N  P  S  Y  V  E  A  Y  N  N    320
CTCCAGATGTACAGAGAAGCCACGCGCCTCAATCCCAGCTACGTCGAGGCTTACAACAAC  960
CTGCAGATGTACCGTGAAGCGACCCGCCTGAACCCGAGCTACGTGGAGGCGTACAACAAC

M  G  A  V  C  K  N  L  G  K  L  E  D  A  I  S  F  Y  E  K    340
ATGGGCGCTGTGTGTAAGAACCTAGGCAAGCTGGAAGACGCTATTTCGTTCTATGAGAAG 1020
ATGGGCGCGGTTTGCAAAAACCTGGGTAAACTGGAAGACGCGATTAGCTTCTACGAGAAA

A  L  A  C  N  A  N  Y  Q  M  S  L  S  N  M  A  V  A  L  T    360
GCCCTTGCATGCAATGCAAACTACCAGATGAGTCTGAGCAACATGGCTGTTGCTCTCACC 1080
GCGCTGGCGTGCAACGCGAACTACCAGATGAGCCTGAGCAACATGGCAGTGGCGCTGACC

D  L  G  T  Q  Q  K  A  S  E  G  A  K  K  A  I  S  L  Y  K    380
GACCTTGGAACTCAGCAGAAAGCTTCTGAAGGCGCGAAGAAGGCAATTTCGCTCTACAAA 1140
GACCTGGGCACGCAGCAGAAGGCAAGCGAGGGTGCGAAGAAAGCGATCAGCCTGTACAAG

K  A  L  I  Y  N  P  Y  Y  S  D  A  Y  Y  N  L  G  V  A  Y    400
AAGGCCTTAATTTACAATCCGTATTATTCGGATGCGTACTACAATCTGGGCGTCGCGTAC 1200
AAAGCGCTGATTTACAACCCGTACTACAGCGACGCGTACTACAACCTGGGCGTTGCGTAC

A  D  L  H  K  F  D  K  A  L  V  N  Y  Q  L  A  V  A  F  N    420
GCCGACTTGCACAAATTCGACAAGGCACTGGTGAACTATCAGCTGGCAGTGGCCTTCAAT 1260
GCGGACCTGCACAAATTCGACAAGGCGCTGGTTAACTACCAGCTGGCAGTTGCATTCAAC

P  R  C  A  E  A  Y  N  N  M  G  V  I  H  K  D  R  E  N  T    440
CCTCGATGTGCTGAGGCGTACAACAACATGGGGGTCATCCATAAGGATAGAGAAAACACG 1320
CCGCGTTGCGCGGAAGCGTACAACAACATGGGTGTTATCCACAAAGACCGCGAGAACACC

D  Q  A  T  V  Y  Y  N  K  A  L  E  I  N  P  D  F  S  Q  T    460
GACCAGGCCACTGTTTACTACAACAAAGCTCTGGAAATAAATCCGGACTTCTCTCAAACG 1380
GACCAGGCGACCGTCTACTACAACAAGGCGCTGGAAATTAACCCGGACTTCAGCCAGACC

L  N  N  L  G  V  L  Y  T  C  T  G  K  I  G  E  A  L  H  F    480
CTCAATAACCTTGGCGTGCTCTACACGTGCACCGGGAAGATTGGCGAGGCGCTGCACTTT 1440
CTGAACAACCTGGGCGTTCTGTACACCTGCACCGGCAAAATCGGCGAGGCGCTGCACTTC

A  K  R  A  I  E  V  N  P  N  Y  A  E  A  Y  N  N  L  G  V    500
GCAAAGCGTGCTATTGAAGTCAATCCGAACTACGCCGAGGCATACAACAACTTGGGAGTT 1500
GCGAAGCGTGCGATTGAAGTCAACCCGAACTACGCGGAGGCGTACAACAACCTGGGCGTG

L  Y  R  D  Q  G  D  I  E  D  S  V  K  A  Y  D  K  C  L  L    520
CTGTACCGGGACCAGGGCGATATCGAGGACTCTGTGAAAGCGTATGACAAATGCTTGCTT 1560
CTGTACCGCGACCAGGGTGACATCGAGGACAGCGTTAAAGCGTACGACAAGTGCCTGCTG

L  D  P  N  S  P  N  A  F  H  N  K  L  L  A  L  N  Y  L  E    540
CTGGATCCCAACTCACCAAACGCCTTCCACAACAAGTTACTGGCGTTGAACTATTTGGAG 1620
CTGGACCCGAACAGCCCGAACGCGTTCCACAACAAACTGCTGGCGCTGAACTACCTGGAA

N  L  P  E  N  E  I  C  R  V  S  E  K  W  G  L  H  F  L  S    560
AATTTGCCTGAAAACGAGATATGTCGCGTTTCTGAGAAGTGGGGACTGCATTTCCTTTCT 1680
AACCTGCCGGAGAACGAAATCTGCCGTGTTAGCGAGAAGTGGGGTCTGCACTTCCTGAGC

S  R  S  P  Y  T  S  W  L  C  P  P  V  T  I  S  P  A  L  P    580
TCTCGGTCTCCGTACACTTCCTGGCTGTGTCCGCCGGTCACCATCTCACCTGCCCTGCCT 1740
AGCCGCAGCCCGTACACCAGCTGGCTGTGCCCGCCGGTTACCATTAGCCCGGCACTGCCG

S  S  A  V  R  S  P  A  R  P  S  S  S  S  A  S  S  S  P  A    600
TCGTCTGCCGTCCGCTCACCTGCCCGTCCATCTTCATCGTCAGCTTCTTCGTCTCCTGCC 1800
AGCAGCGCGGTCCGTAGCCCGGCGCGCCCGAGCAGCAGCAGCGCAAGCAGCAGCCCGGCA

S  P  G  D  S  S  A  S  R  V  I  R  V  G  Y  I  G  P  D  F    620 D619A TCTCCTGGCGATTCTTCAGCTTCAAGAGTCATACGCGTGGGGTACATTGGACCAGATTTC 1860
AGCCCGGGTGACAGCAGCGCGAGCCGTGTGATCCGCGTTGGCTACATTGGTCCGGACTTC

F  T  H  S  V  S  Y  F  I  H  A  P  L  V  Y  H  D  K  A  K    640 H623A TTTACTCATTCTGTCTCTTACTTCATCCATGCTCCCTTGGTCTACCACGACAAAGCGAAG 1920
TTCACCCACAGCGTCAGCTACTTCATCCACGCGCCGCTGGTGTACCACGACAAGGCGAAA

F  H  I  T  V  Y  A  N  V  I  R  E  D  E  K  T  Q  M  F  K    660
TTCCACATCACTGTCTACGCGAACGTCATCCGAGAAGACGAAAAGACTCAAATGTTCAAG 1980
TTCCACATCACCGTTTACGCGAACGTCATTCGTGAGGACGAAAAAACCCAGATGTTCAAG

T  L  P  H  R  W  R  S  I  V  G  L  N  E  Q  E  V  A  R  I    680
ACGCTCCCGCACCGCTGGCGGTCTATCGTGGGGTTGAACGAGCAGGAGGTTGCTCGGATC 2040
ACCCTGCCGCACCGTTGGCGCAGCATTGTCGGCCTGAACGAGCAGGAAGTGGCGCGTATC

I  R  E  E  D  R  I  D  I  L  V  E  L  A  G  H  T  A  H  N    700 E692K,G695D ATCCGAGAAGAAGACCGAATCGACATTTTGGTGGAACTCGCAGGGCACACAGCGCACAAC 2100
ATTCGCGAGGAAGACCGTATCGACATTCTGGTCGAACTGGCAGGTCACACCGCACACAAC

R  L  D  V  M  A  C  K  P  A  P  V  Q  I  S  W  I  G  Y  P    720
CGCCTCGACGTGATGGCGTGCAAACCTGCGCCGGTTCAGATCAGCTGGATTGGCTATCCG 2160
CGTCTGGACGTTATGGCGTGCAAACCGGCGCCGGTCCAGATCAGCTGGATTGGCTACCCG

N  T  T  G  L  K  T  I  D  F  R  I  T  D  A  V  A  D  P  L    740
AACACAACCGGCTTGAAGACCATCGACTTCCGCATCACTGACGCCGTCGCGGACCCGCTG 2220
AACACCACCGGTCTGAAGACCATCGACTTCCGTATTACCGACGCAGTGGCGGACCCGCTG

T  T  T  E  R  Y  V  E  E  L  V  R  M  P  N  C  F  L  C  Y    760
ACCACAACGGAGAGGTATGTGGAAGAGCTGGTCCGCATGCCCAACTGCTTCCTCTGCTAC 2280
ACCACCACCGAGCGTTACGTGGAGGAACTGGTTCGCATGCCGAACTGCTTCCTGTGCTAC

Q  P  P  P  D  F  P  K  H  V  P  A  K  P  P  P  V  L  D  H    780
CAGCCGCCTCCCGACTTCCCGAAGCATGTGCCAGCGAAGCCGCCCCCCGTTCTCGATCAC 2340
CAGCCGCCGCCGGACTTCCCGAAACACGTTCCGGCAAAGCCGCCGCCGGTCCTGGACCAC

G  V  V  T  F  G  S  F  N  N  L  A  K  L  G  N  Q  V  I  E    800 K793A  GGCGTCGTCACCTTCGGCTCGTTCAACAATCTCGCAAAACTCGGCAATCAGGTGATTGAG 2400
GGCGTGGTTACCTTCGGTAGCTTCAACAACCTGGCGAAACTGGGTAACCAGGTGATCGAA

I  W  S  R  I  L  N  A  V  P  N  S  R  L  L  V  K  A  R  P    820
ATCTGGAGTCGGATTTTGAATGCCGTCCCGAATAGCCGGTTGCTGGTCAAAGCCCGGCCG 2460
ATTTGGAGCCGCATCCTGAACGCGGTGCCGAACAGCCGTCTGCTGGTTAAAGCGCGCCCG

F  A  N  K  E  M  Q  R  K  F  K  A  K  F  E  A  H  G  I  S    840
TTTGCAAACAAGGAAATGCAGCGGAAATTCAAGGCGAAATTTGAAGCGCATGGAATCTCG 2520
TTCGCGAACAAGGAAATGCAGCGTAAGTTCAAAGCGAAGTTCGAGGCGCACGGCATCAGC

G  D  R  I  D  A  M  A  L  I  P  A  C  M  D  H  L  M  V  Y    860
GGCGATCGTATCGACGCCATGGCTCTCATTCCTGCATGCATGGATCACTTAATGGTCTAT 2580
GGTGACCGCATTGACGCGATGGCGCTGATCCCGGCGTGCATGGACCACCTGATGGTCTAC

S  L  V  D  I  A  L  D  S  F  P  Y  A  G  T  T  T  T  C  E    880
TCGCTAGTTGACATCGCCCTCGATTCCTTCCCTTACGCCGGAACAACAACGACCTGTGAG 2640
AGCCTGGTGGACATTGCACTGGACAGCTTCCCGTACGCGGGTACCACCACCACCTGCGAG

A  L  V  M  G  V  P  V  V  S  L  R  R  P  N  I  H  A  H  N    900
GCACTTGTCATGGGCGTCCCTGTCGTCTCTCTTCGTCGCCCAAACATCCACGCACACAAT 2700
GCACTGGTCATGGGTGTGCCGGTTGTGAGCCTGCGTCGCCCGAACATTCACGCACACAAC

V  G  A  T  L  L  V  N  Y  G  L  P  E  L  I  A  D  D  P  E    920
GTAGGAGCAACTCTGCTGGTTAACTACGGACTGCCCGAACTTATCGCAGACGATCCCGAA 2760
GTGGGTGCAACCCTGCTGGTTAACTACGGTCTGCCGGAACTGATTGCGGACGACCCGGAG

Q  Y  V  R  V  A  V  E  L  A  G  D  V  E  R  L  K  R  Y  R    940
CAATATGTTCGTGTAGCTGTCGAGCTCGCAGGGGATGTCGAGCGCCTAAAACGCTATCGG 2820
CAGTACGTTCGTGTTGCAGTGGAACTGGCAGGTGACGTGGAGCGTCTGAAGCGTTACCGC

Q  S  I  R  E  S  V  L  E  K  A  S  E  P  H  A  K  Q  F  T    960
CAGAGCATCCGCGAATCAGTTCTCGAGAAGGCGTCTGAGCCGCATGCGAAGCAGTTCACT 2880
CAGAGCATCCGCGAGAGCGTTCTGGAAAAAGCGAGCGAGCCGCACGCGAAGCAGTTCACC

R  D  L  E  E  L  Y  R  Q  L  L  A  R  K  H  R  Q  K  P  R    980
CGCGACTTAGAGGAGCTGTATCGCCAGCTGCTAGCTCGCAAACACCGGCAAAAGTAG    2937
CGTGACCTGGAGGAACTGTACCGCCAGCTGCTGGCGCGTAAACACCGCCAGAAGCCTAGG

*
TAAggatcctaataactaagtaaactagtgctgagcaataactagcataaccccttgggg

cctctaaaggcccggcagtaccggcataaccaagcctatgccTacagcatccagggtgac

ggtgccgaggatgacgatgagcgcattgttagatttcatacacggtgcctgactgcgtta

gcaatttaactgtgataaactaccgcattaaagcttatcgatgataagctgtcaaacatg

agaa
```

**Figure S2. Genotyping of *T. gondii* genetically modified strained described in this study.** *A*. PCR analysis verifying the absence of endogenous *spy* (SPYe) in the knockout and complemented strains. Additionally, the DHFR cassette has been removed from the *spy* locus in the knockout strain used here. Codon reassigned SPY (SPYc) was used to complement the knockout strains and can be selectively amplified by PCR. *B*. Schematic representation of the strategy used to knockout *Tg*SPY in the type II strain CZ1. *C*. Southern blot analysis was used to verify single integration of the mGFP cassette in the *spy* locus of CZ1 ∆*ku80* *∆hpt* Re9. *D*. Schematic representation of the domains of *Tg*GPN. Residues 1-79 were defined as SRD and *Tg*GPN∆SRD starts at aa 80. The residues marked with asterisks are part of the active site of GPNs (Ref. S1, S2). *E*. Genotyping by PCR of the clones expressing either full length or truncated GPN. All primers are listed in Table S1. Pf: *Plasmodium falciparum*, Pp: *Pyrococcus abyssi*.

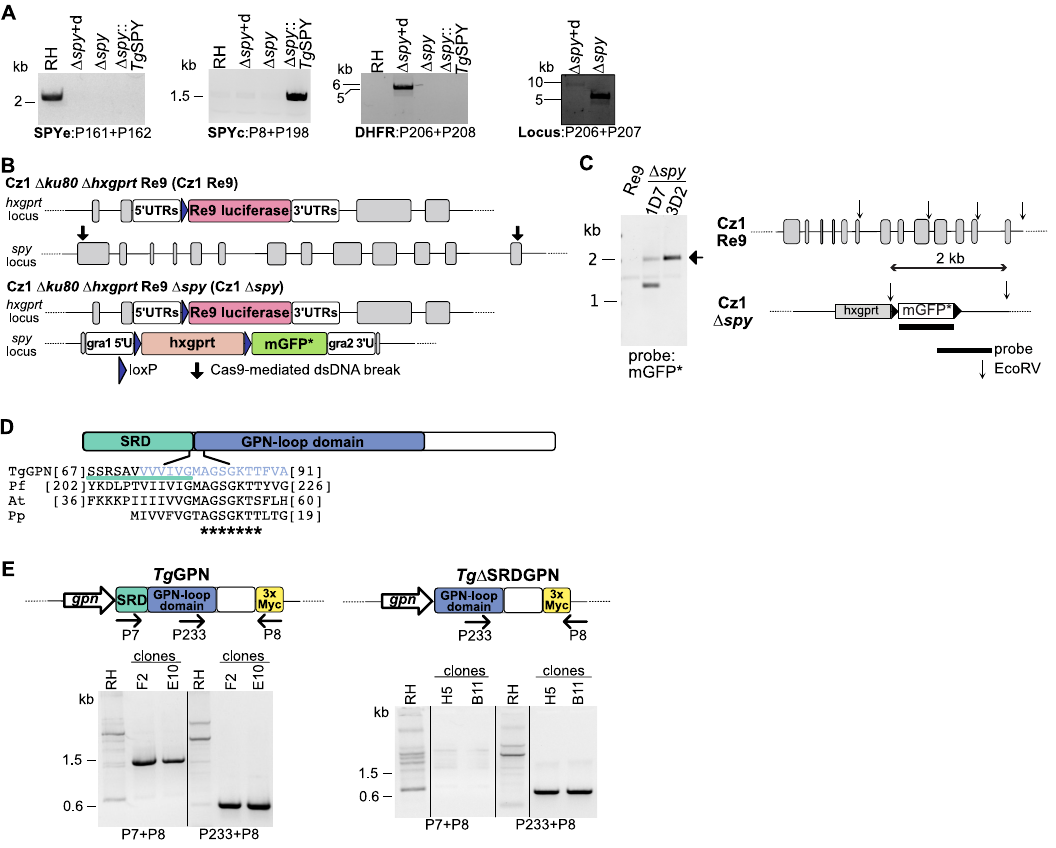

**Figure S3. Phenotype scores indicate *O*-fucosylated proteins are important for tachyzoites fitness.** Violin plots comparing the mean phenotype score distributions of the complete set of AAL enriched proteins (n=69), the subset for which glycopeptides were identified (n=33) and the subset composed of hypothetical conserved proteins (n=29) with known dispensable and essential genes for tachyzoites survival in culture (text ref. 50). Violin plots were generated using R. The white dot represents the median and the thick black line the interquartile range.

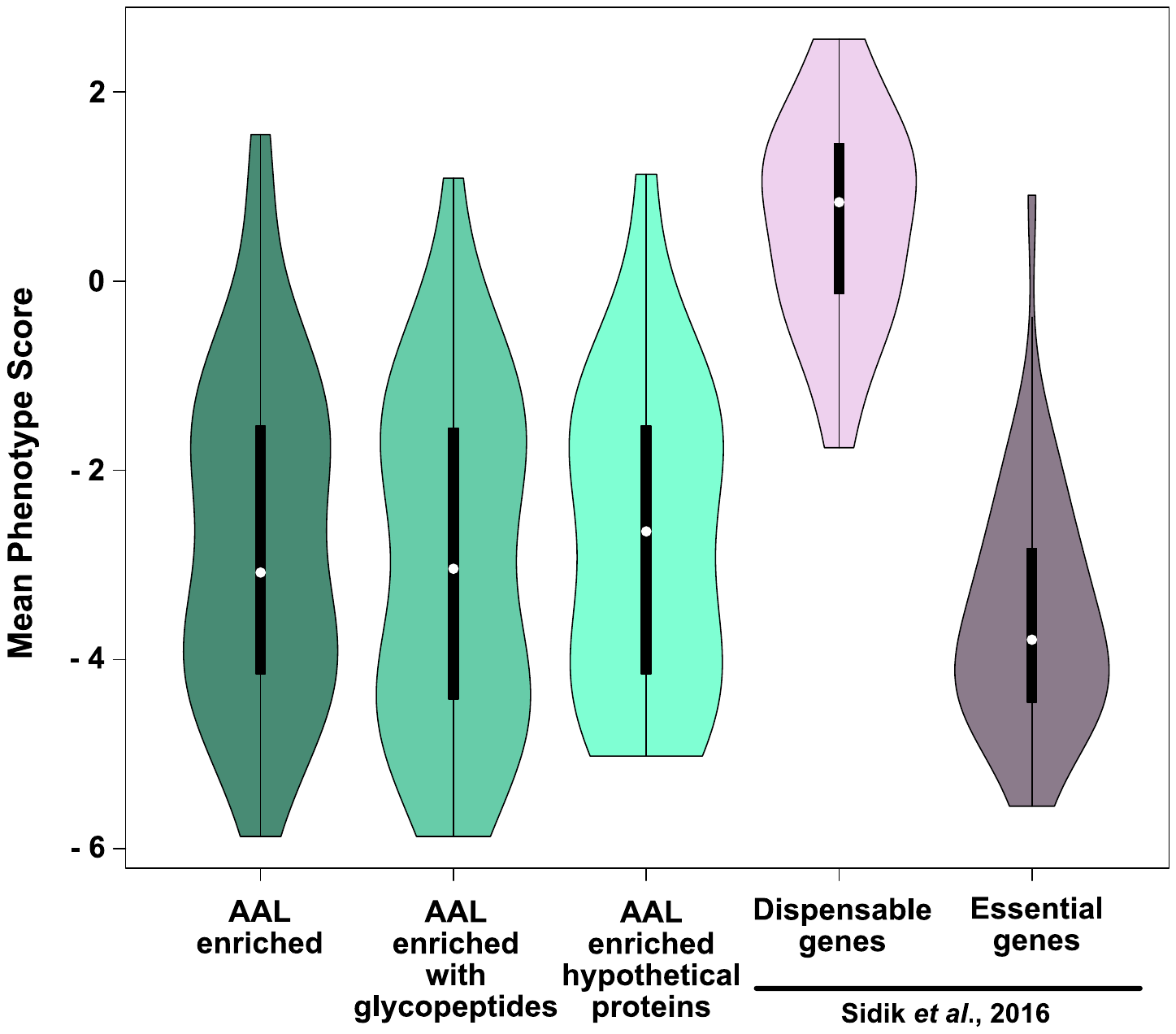
